# Single-cell dissection of bacterial communities into functional gene carriers with ddPCR

**DOI:** 10.1101/2022.09.27.509627

**Authors:** Lara Ambrosio Leal Dutra, Matti Jalasvuori, Ole Franz, Kimi Nurminen, Pauliina Salmi, Marja Tiirola, Reetta Penttinen

**Affiliations:** Department of Biological and Environmental Science and Nanoscience Center, University of Jyväskylä, Jyväskylä, Finland; Department of Biology, University of Turku, Turku, Finland

**Author notes:** Correspondence to Reetta Penttinen,.

**Keywords:** 16S rRNA gene, droplet digital PCR, droplet microfluidics, functional genes, microbial diversity analysis, single-cell

## Abstract

Microbial communities often respond to environmental challenges, such as the presence of antibiotics, as a whole. Dissecting these community-level effects into separate acting entities requires the identification of organisms that carry functional genes for the observed feature. However, unculturable microbes are abundant in various environments, hence making the identification challenging. Moreover, while at present the development and application of single-cell tools for eukaryotic cells are enhancing, the comparable methodologies applicable for prokaryotic cells are still scarce and have not gained broad and solid status as tools for investigating microbial populations. Here, we present a cultivation-free technique that can be utilized to link functional genes with the carrying bacterial species at single-cell resolution. The developed protocol is relatively simple to use, utilizes commercially available droplet microfluidics devices, does not require toxic reagents, and eliminates invalid signals emerging from extracellular DNA. We validate the methodology by studying the conjugative transfer of antibiotic resistance plasmids in an environment challenged by antibiotics. Furthermore, the method can be customized for any given genetic trait to accurately identify its hosting subpopulation from a heterogeneous and potentially uncultivable bacterial community.

**Importance:** Bacterial systems usually contain numerous different species that may harbor highly similar or identical genes that confer same phenotypic qualities for the community. To decipher the functions of these systems, we report the development of a novel methodology that enables investigating microbial communities at single-cell level. This user-friendly method utilizes droplet digital PCR (ddPCR) to find and identify carriers of specific genes potentially from various microbial sample types. By pinpointing gene carriers, such as those responsible for antibiotic resistance, this method can provide insights to the behavior of microbial communities and gene transfer therein. By strategically combining the use of common methods (ddPCR and amplicon sequencing), the workflow is highly accessible. Thus, it allows also the researchers without a background in single-cell techniques or access to special equipment to adopt the method for producing single-cell data to serve their own research, enabling new research avenues in microbial genetics and ecology.

## Introduction

Microbes are ubiquitous organisms that play important roles in systems shaping life on our planet. Yet, microbes rarely exist in isolation. A single grain of sand contains tens to hundreds of thousands of bacterial species^1^, one gram of dry faecal matter has a billion microbial cells and even very slowly growing microbial mats in lava caves have hundreds of distinct taxonomical units in their thick, slimy layers^2^. Within these communities, bacteria form interaction networks where individual cells are often unable to reproduce in the absence of their contemporaries^3^. Ultimately, genetic, metabolic and ecological links between microbes shape the communities and their behaviour^4^. Bacterial systems may, for example, degrade hazardous waste materials, confer resistance to antibiotics and regulate the turnover of limiting nutrients for subsequent levels in trophic cascades. Further, horizontal gene transfer is common in microbial systems and opportunistic genetic characteristics may rapidly switch into more suitable hosts in response to environmental conditions^5^. Disentangling these communities into individual cells that can still be studied with precision and without the influence of potential confounding factors, such as remnant DNA from dead cells, is challenging.

Various methodological advances have been achieved in the recent years to help resolve the roles of individual and potentially unculturable microbes within communities. 16S ribosomal RNA sequencing is the current standard for providing an overall picture of the diversity of microbes – living or dead – in the studied sample. However, it fails to associate genetic characteristics to the species. In contrast, total DNA sequencing can cover a lot of the overall genetic variation, but usually without a definitive answer to the question of who acted as the cellular vehicle for any particular gene – or if that gene was indeed actually a part of a living organism at all or merely a remnant from a deceased microbe^6^. Further, (mobile) plasmids in particular, replicate separately from chromosomes in bacterial cells, and hence, metagenomic sequencing fails to identify the host of the potentially crucial genetic element^7^. To link genetic traits with particular species, a number of microfluidic techniques have been utilized to identify target traits (such as the ability to cause a disease or carry bacteriophages) from heterogeneous bacterial samples^8,9^. Ottesen and colleagues utilized a chip containing 1176 microchambers to divide termite midgut samples to single-cell resolution and used two fluorescent probes to determine the chambers containing 16S rRNA and/or the targeted functional gene^10^. By using a needle to extract the contents of chambers that expressed both fluorescent signals, they were able to associate 16S rRNA sequence with the functional gene. Alternatively, fixed bacterial samples have been used for fluorescent probe hybridization^11^. Hybridization methods, however, are relatively laborious and unable to study a large number of cells. In order to study hundreds of thousands of cells from uncharacterized bacterial samples, Spencer and colleagues developed a method to trap individual cells into tiny acrylamide compartments prior to PCR amplification^12^. In this so-called epicPCR, a very localized PCR reaction is utilized to amplify the targeted functional gene and fuse it in concatenation PCR with the 16S ribosomal RNA gene. By sequencing the paired products, the cellular hosts harbouring the studied genetic trait can be identified. While epicPCR is a promising tool for large-scale, single-cell resolution analyses, it is associated with sources of variation that need to be controlled, including the quantification of the template and cross-contamination.

Droplet microfluidics utilizes microchannels for the formation of nanolitre-scale droplets. In droplet digital PCR (ddPCR), PCR reagents are enclosed into droplets surrounded and stabilized by oil, enabling thousands of PCR reactions to occur simultaneously. The amplified DNA is detected by using double-stranded DNA-binding fluorescent dye and measured from each droplet one by one. As the sample is randomly distributed across the droplets, both empty (negative) and DNA-containing (positive) droplets are formed. The system is extremely sensitive and capable of detecting even a single copy within a droplet. Finally, the fraction of positive droplets is compared to the total number of droplets and used to calculate the (absolute) quantity of target DNA in the original sample with the help of Poisson statistics^13,14^. DNA can be delivered into droplets also within intact cells instead of pure DNA, allowing ddPCR to be applied for bacterial communities. This principle has been harnessed to trace enterohemorrhagic *Escherichia coli* (EHEC) in food production samples^15,16^. In these studies, single bacterial cells were enclosed into droplets. After the cell lysis initiated by the denaturation step of ddPCR, EHEC was detectable via distinct amplification of two virulence genes and their colocalization in the same droplet with fluorescent probes^16^.

In this study, we present a new adapted technique for investigating bacterial communities at single-cell resolution. The described method utilizes commercially available ddPCR devices (based on droplet microfluidics), is fast to perform, requires only non-toxic reagents, and contains steps to efficiently inactivate extracellular DNA from samples. The method employs paired fusion PCR to link a taxonomic marker to the target gene and allows accurate fluorescence-based verification of single-cell resolution prior to fusion amplicon sequencing. We demonstrate that the method is applicable for precise detection and identification of carriers of a beta-lactamase resistance gene (*bla*CTX-M-14) in the heterogenous bacterial sample, and thus, can be utilized for tracing the movements of mobile genetic elements such as plasmids within the bacterial community. As such, the introduced single-cell ddPCR method provides a convenient approach for different laboratories to identify specific genetic traits from multi-species communities.

## RESULTS

### A microbial single-cell screening from a community

In this study, we developed an easy-to-use microbial single-cell screening method to identify or track hosts of any gene of interest in heterogeneous communities. Our experimental setup focused on tracking an antibiotic resistance gene (ARG) *bla*CTX-M-14 (hereafter referred to as *bla*). A fragment of *bla* is paired with the 16S rRNA gene in fusion PCR to identify taxonomic groups that harbour the ARG. For enabling analysis of individual cells, the bacterial samples are partitioned into compartments by utilizing the QX200™ Droplet Digital PCR System from Bio-Rad. This automated system employs droplet microfluidics for generating equal-sized 1-nanolitre droplets that encapsulate both individual bacterial cells and PCR reagents (Figure 1). During thermal cycling, the bacterial DNA is released from cells via lysis and used as a template for paired amplification of target ARG and 16S rRNA gene. As single-cell PCR-based methods are highly sensitive to any contaminating DNA present in the droplet but outside of the cell, the protocol includes the treatment of the sample prior partitioning for inactivating extracellular DNA. These steps were observed to be crucial for the accurate detection of genetic content that is indeed bound within cellular membranes. By sequencing the paired amplicons, the *bla* carrier population can be identified from the studied bacterial community. Four bacterial species were employed in the development and optimization of the described method: *Escherichia coli* (carrying the target *bla* gene in a conjugative plasmid pEC13), *Pseudomonas fluorescens, Salmonella* Typhimurium and *Klebsiella pneumoniae*.

**Figure 1.**
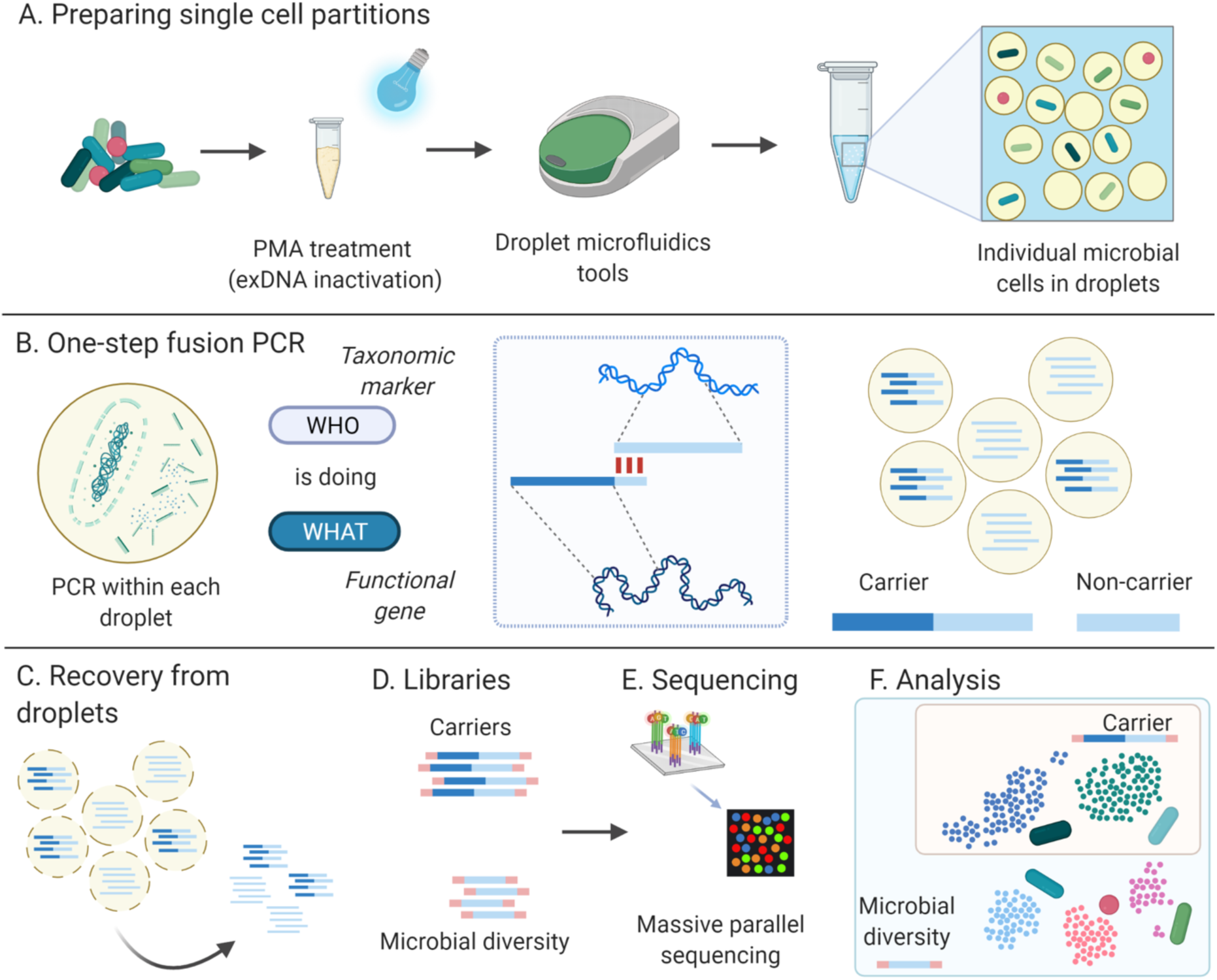
Overview of the single-cell ddPCR method workflow. A) Bacterial samples are prepared and partitioned using droplet microfluidics technology to obtain a single cell per droplet. This step includes treatment with PMA (propidium monoazide) to inactivate exDNA (extracellular DNA), the addition of PCR reagents and, finally, the generation of single-cell - containing droplets with a droplet microfluidics device. B) One-step fusion PCR assay is carried out within each droplet. In droplets containing a carrier cell, *i.e.* a cell that has the functional gene, the taxonomic marker gene and the functional gene are amplified and fused into a concatemer PCR product. The taxonomic marker is the sole product in droplets containing non-carrier bacterial cells. C) The produced amplicons are recovered from droplets by breaking the emulsion using a liquid nitrogen method^17^. D) The recovered amplicons are used to construct sequencing libraries and E) sequenced by massive parallel sequencing. F) The sequence analysis determines the bacterial population carrying the functional gene as well as the overall bacterial diversity in the given sample. Created with BioRender.com.

### Extracellular DNA (exDNA) inactivation from cell samples

Microfluidic encapsulation traps any content in the sample, including free DNA. Preliminary analysis (not shown) indicated that free DNA (*i.e.* DNA outside of bacterial cells) is abundant in bacterial suspensions even after DNAse treatment and/or repeated washing of the cells. As such, this DNA forms a notable confounding factor for single-cell analyses that aim to reliably identify a specific DNA target that is indeed within the cellular membranes (*i.e.* a single viable cell). We ensured that the process only amplifies DNA originating from a unique cell by treating the samples with propidium monoazide (PMA). PMA is a DNA binding dye that covalently attaches to double stranded DNA (dsDNA) via photoactivation, thus preventing primer annealing and polymerization to occur. Cell membranes are impermeable to PMA, hence only DNA external to intact cells is rendered non-amplifiable. Phosphate buffer saline (PBS) was used for diluting the bacterial cell samples, therefore its maximum non-inhibitory concentration in ddPCR was defined and always kept at safe level (under 0.0625x) (see Supplementary Figure S1).

### PMA treatment – fluorescence emission spectral shift

After initial screening, we hypothesized that PMA may influence PCR. At first, we wanted to see if PMA would prevent amplification, therefore we expected the number of positive droplets would reduce with increasing amounts of PMA. As such, the maximum non-inhibitory concentration of PMA in PCR amplification was assessed by adding increasing concentrations of photoactivated PMA to EvaGreen-based ddPCR assays (Figure 2). The reactions targeting the 16S rRNA region (amplified with 27F and 338R primers) of digested *E. coli* DNA were prepared with 0.125 µM up to 2.5 µM of PMA. The Relative Fluorescence Unit (RFU) was obtained from each droplet by the Droplet Reader through fluorescence detection channels 1 (Ch1 for FAM) and 2 (Ch2 for HEX/VIC). Analysis of RFU showed that two droplet clusters could be seen: a cluster of negative droplets where amplification had not occurred (low RFU) and a cluster of positive droplets with successful amplification (high RFU). Nonetheless, in Ch1, a pronounced drop in RFU from positive droplet cluster could be seen when increasing concentrations of PMA were used (Figure 2A). The same pattern could not be observed in Ch2 data. Here, instead, the positive droplets cluster showed an increase in RFU in higher PMA concentrations. The analysis of Ch1 *versus* Ch2 (Figure 2B) showed a clear distinction between positive and negative droplet clusters. These results suggest that PMA does not inhibit PCR reactions when used in concentrations up to 2.5 µM but affects the fluorescence emission spectrum that disturbs the detection of the DNA binding dye (EvaGreen) from Ch1.

**Figure 2.**
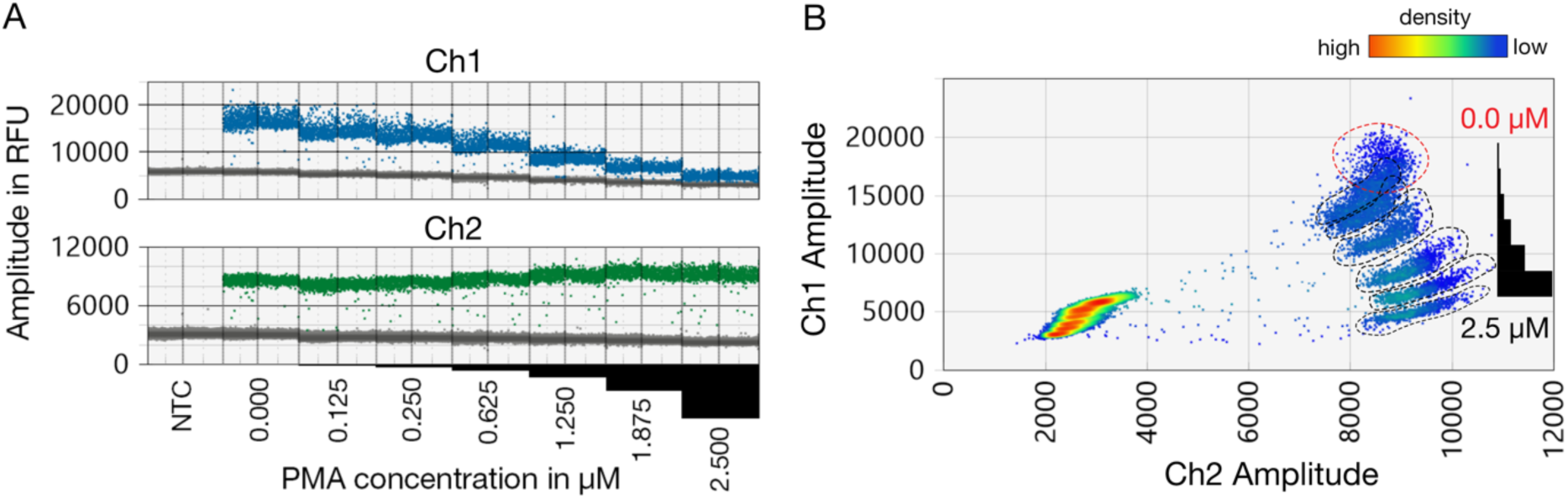
Effect of PMA concentration on RFU reading. Light-inactivated PMA in final concentration ranging from 0.125 to 2.5 µM was added to 16S rRNA ddPCR. After amplification, the fluorescence of each droplet was measured through both detection channels. The increasing PMA concentration resulted in a drop in RFU when measured via channel 1 (Ch1 for FAM) and 2 (Ch2 for HEX/VIC).) (A, upper panel), but not with channel 2 (Ch2) (A, lower panel). The positive droplets were identified by two-dimensional plotting of RFU values detected from Ch1 and Ch2 (B). The circles mark roughly the cluster of positive droplets for the different PMA concentration.

Further, we studied whether PMA inactivates exDNA in samples containing bacterial cells. Cell suspensions of *E. coli* and *P. fluorescens* and their filtrates (suspension filtered through a 0.2 µm filter) were treated with 2 µM PMA and used as a template in ddPCR amplifying a region within the gene for 16S rRNA. For controls, the corresponding samples were treated with sterile water instead of PMA. PMA treatment reduced the obtained 16S rRNA concentration by as much as 71% in *P. fluorescens* and 22% in *E. coli* cell samples (Figure 3). This suggests that exDNA is abundant even in pure bacterial cultures and potentially causes erroneous PCR amplification if the DNA is present inside a droplet together with the trapped bacterial cell. To verify the complete inactivation of exDNA, we filtered out the bacterial cells and exposed the filtrate to PMA. Less than 0.12% of the original contaminating DNA was amplifiable after PMA treatment (Figure 3).

**Figure 3.**
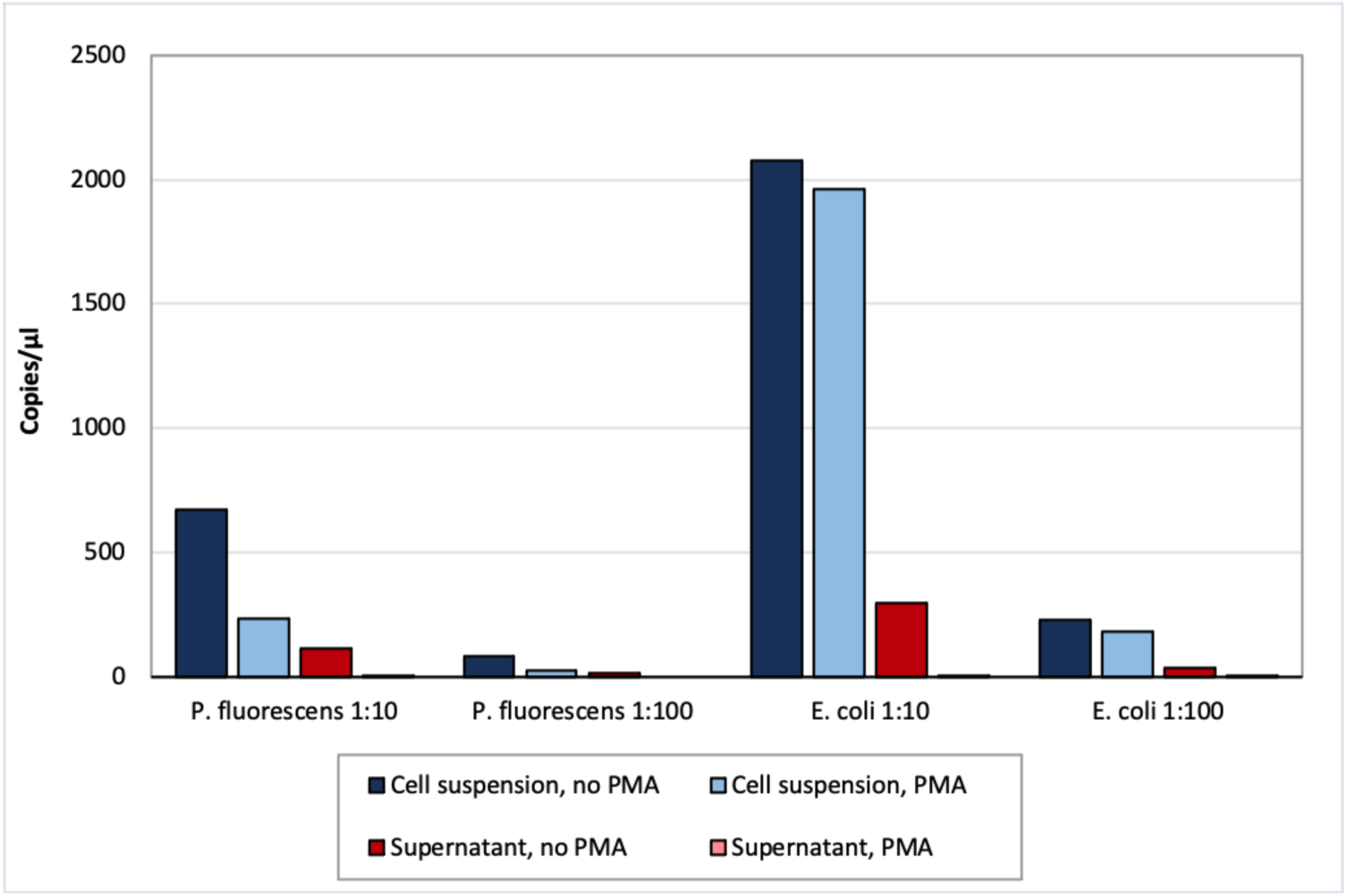
Effect of PMA treatment on inactivation of exDNA. *Pseudomonas fluorescens* and *Escherichia coli* cell dilution (1:10 or 1:100 of an overnight-grown bacterial culture) and their filtrated (0.2 µm) supernatant were treated with 10 µM PMA or sterile water as control. 16S rRNA was amplified with single-cell ddPCR and its abundance is presented as copies/µl.

### One-step fusion PCR

The developed method relies on linking two targeted DNA fragments together by using fusion PCR. To produce the concatemer, we employed one-step fusion PCR with four primers. In this approach, the amplification of both targets occurs within the same reaction. Introduction of a fusion primer into the reaction prompts the attachment of the distinct target fragments into a unified DNA molecule (Figure 4). The fusion primer consists of a priming sequence for a given target sequence added by additional bases at the 5’-end that encode the complementary reverse sequence of the forward primer of the other target. The dynamics of the reaction is driven by a difference in the ratio of forward to reverse primer (within a pair) and the concentration of each primer pair. The outer primers that flank the concatemer sequence are used in higher concentrations than the inner primers. Amplification of the target sequence will eventually diminish due to the depletion of inner primers, thus the matching overhang will prompt the fusion of both target sequences into a paired concatemer. The excess of concatemer flanking primers allows for the amplification of the linked sequence.

**Figure 4.**
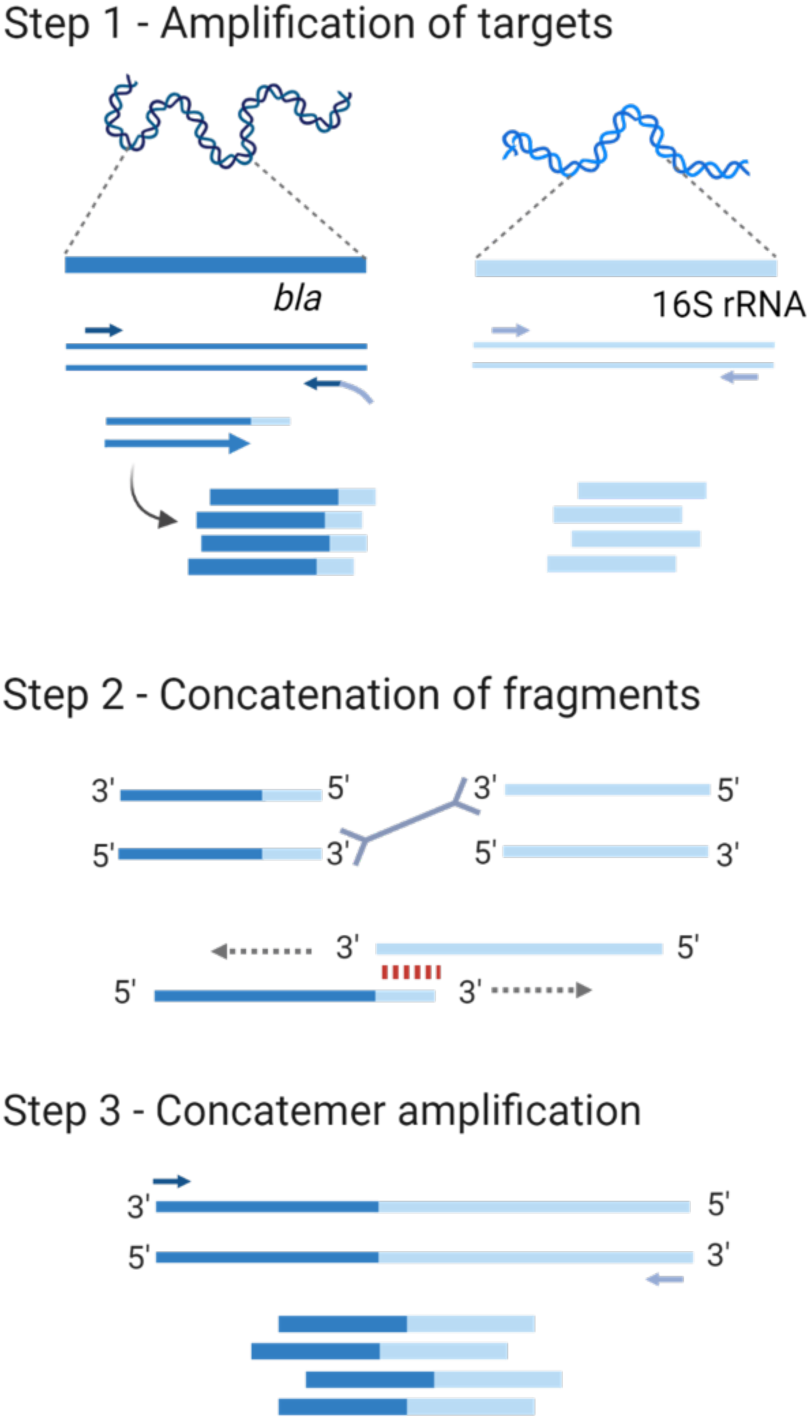
One-step fusion PCR with four primers. The amplification and fusion of 16S rRNA and beta-lactamase gene *bla* are carried out in the same reaction. The fusion process is driven by a fusion primer that is one of the *bla* primers with a 5’ -end extension that encodes the reverse complement sequence of the 16S rRNA primer 27F. In step 1, both target regions are simultaneously and independently amplified, generating the intermediate product of the fusion: the *bla* fragment with the tail complementary to 27F. In step 2, the tail will prompt the fusion of 16S and *bla* products. Lower concentration of the inner primers (27F and the modified *bla* primers that bind to the fusion region) facilitates the reaction dynamic to produce the concatemer. After the depletion of the inner primers, in step 3, the amplification of the fusion product is carried out by using the excess of external primers. Created with BioRender.com.

In our experiments, we used the 16S rRNA universal primers, 27F and 338R, to amplify a variable region that can be used to identify the used bacterial strains. As for ARG target, we designed primers 293F and 320R for amplifying a 101 bp region of the *bla* gene. Furthermore, we modified *bla* reverse primer 320R with a 5’-extension of the reverse complementary sequence of 27F (therefore producing a fragment of 121 bp). The complementary sequence to be produced during PCR was to act as primer and allow annealing of the *bla-*27F overhang to the 16S rRNA fragment, thus fusing both target sequences as a concatemer.

### Optimization of primer sets

The optimization of one-step fusion PCR was carried out by screening the primer ratio and concentration to assess the efficiency of concatemer production. First, the pairs were tested separately to verify the production of each target in different reaction conditions (Supplementary Figures S2-3). After this, the primers were combined in a single ddPCR reaction to ensure the amplification of the concatemer. The separation of positive and negative clusters was evaluated to define the concentration and ratio to be used in the following assays. Clustering was more efficient at lower ratios and with higher primer concentrations (Supplementary Figure S3).

In ddPCR, the amplification of DNA fragments of distinct sizes is indicated by RFU level. Thus, the occurrence and separation of four droplet clusters were sought: a cluster with very low RFU was expected for negative droplets; two clusters with intermediate RFU were expected to result from droplets where a single target was amplified, either a low RFU cluster for *bla* (121 bp product) or higher RFU cluster for 16S rRNA (350 bp product). The cluster with the highest RFU indicated the amplification of the fused concatemer (of 450 bp) (for detailed information, see Supplementary Figure S3). Finally, the conditions for one-step fusion reactions in ddPCR were defined by using 225 nM of 16S rRNA primers in 1:5 (forward:reverse) ratio, and 200 nM of *bla* primers in 10:1 (forward:reverse) ratio.

### Partition of single cells into droplets

In order to verify whether single cells could be delivered to droplets with the ddPCR microfluidic system, we designed an assay where PMA-treated dilutions of *E. coli, Pseudomonas fluorescens, Salmonella* Typhimurium and *Klebsiella pneumoniae* cells were used as a template in the ddPCR targeting the 16S rRNA gene. A serial plating assay was prepared in parallel, and the obtained cell density compared to that estimated with ddPCR. With all tested bacterial strains, the bacterial cell density was comparable when measured with serial plating and 16S rRNA single-cell ddPCR; the decreasing trend was detectable between adjacent dilutions. However, in *P. fluorescens* and *K. pneumoniae*, a higher cell concentration was consistently observed with ddPCR in comparison to plating (Figure 5).

**Figure 5.**
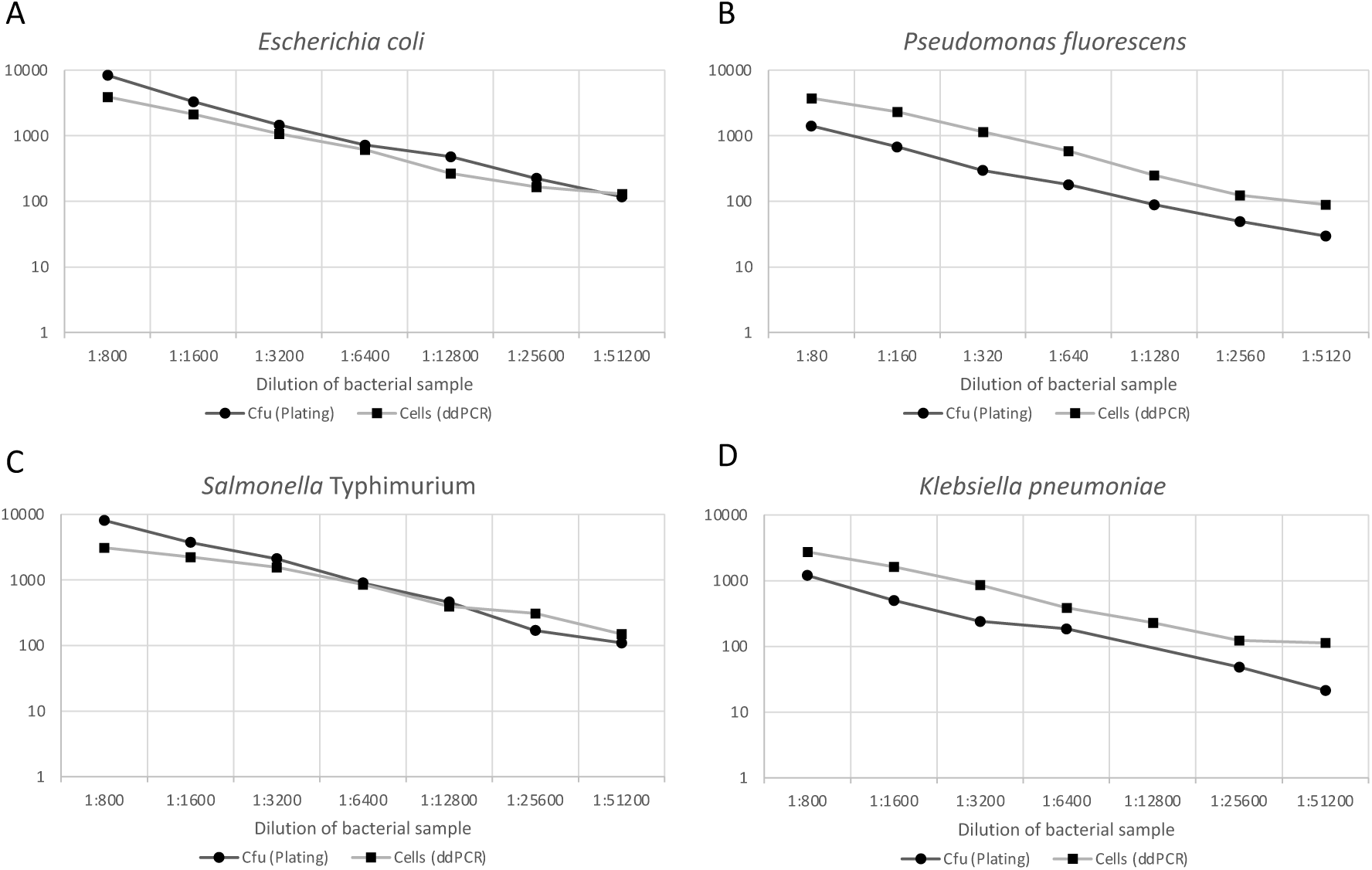
Single cells of *Escherichia coli, Pseudomonas fluorescens, Salmonella* Typhimurium and *Klebsiella pneumoniae* within droplets. A dilution series of overnight-grown bacterial cultures was prepared, enclosed into droplets and amplified with 16S rRNA single-cell ddPCR. The cell number per ddPCR sample was determined by 16S rRNA ddPCR and is presented as cells. Parallel serial plating of bacterial cultures on LB agar was performed to measure cfu (colony forming units) per sample.

### Identifying the ARG carriers by concatemer sequencing

To investigate how effectively the single-cell ddPCR method identifies target gene hosts from a heterogeneous sample, *E. coli* cells carrying *bla*CTX-M-14 were mixed with *S.* Typhimurium and *K. pneumoniae* cells not carrying the gene. Bacterial cell suspensions were first pre-treated with PMA and immediately added to ddPCR reactions to include either a single strain or two strains: *E. coli* with either *S.* Typhimurium or *K. pneumoniae.* The portion of each strain was determined by i) serial plating of each bacterial pure culture, ii) single-cell ddPCR amplifying 16S rRNA and iii) OTU abundance (determined via sequencing of bla_16S concatemer produced in single-cell ddPCR) (Figure 6). The portion of *E. coli* cells in the samples ranged from 28-61% when estimated by separate serial plating of the pure cultures. A comparable ratio was obtained with ddPCR from a mix of *E. coli* and S. Typhimurium cells, however from *E. coli* and *K. pneumoniae* cell mixes, ddPCR resulted in higher portion of *K. pneumoniae* cells. The mixed cell samples were delivered into single-cell ddPCR in which bla_16S concatemers were produced and proceeded to library preparation and sequencing. The 16S rRNA part of the concatemer was used for identifying the host strain associated with *bla*CTX-M-14 and the abundance of *E. coli*, *S.* Typhimurium or *K. pneumoniae* hosts was determined in each sample. In the mixed samples, the portion of non-carrier (*S.* Typhimurium or *K. pneumoniae*) hosts was 0.47-1.15%, suggesting that single-cell ddPCR process effectively diminished the occurrence of false positive host sequences.

**Figure 6.**
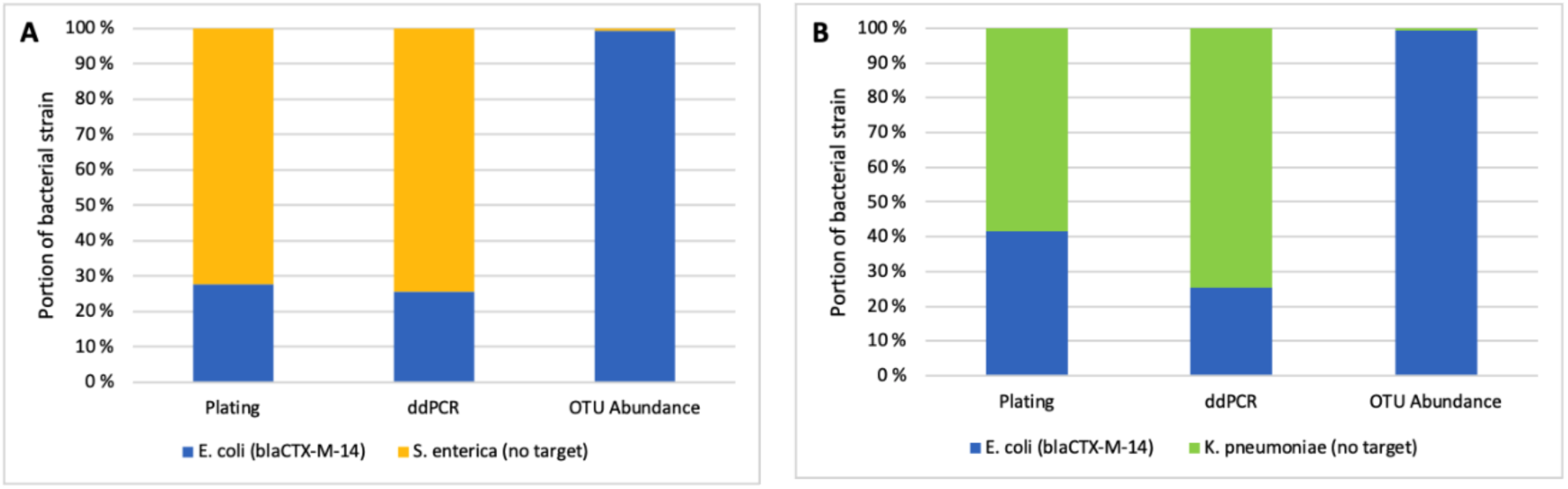
Portions of *Escherichia coli, Salmonella* Typhimurium (A) and *Klebsiella pneumoniae* (B) cells in mixed heterogeneous samples determined by serial plating, 16S rRNA single-cell ddPCR and bla_16S concatemer single-cell ddPCR and sequencing. Cell density (colony forming units/ml) from each pure culture was measured separately by serial dilution and plate counting. 16S rRNA single-cell ddPCR was used to measure the cell density of each pure culture separately to determine the portion of each bacterial strain in a mixed sample. Single-cell ddPCR with primers producing bla_16S concatemer followed by fusion amplicon sequencing was utilized to identify and quantify the *bla*CTX-M-14 carriers in mixed samples. The fraction of *bla*CTX-M-14 hosts is presented as OTU abundance.

### Horizontal transfer of beta-lactamase *bla*CTX-M-14 in synthetic bacterial community

The developed method was applied to track horizontal transfer of a conjugative plasmid within a synthetic microbial community consisting of rifampicin resistant *Klebsiella pneumoniae, Comamonas* sp. and *Salmonella* Typhimurium strains. As a plasmid donor, we harnessed rifampicin-sensitive *E. coli* strain with conjugative plasmid pEC13, which renders the host resistance to cephalothin via *bla*CTX-M-14 gene. This plasmid has been shown to be able to rescue antibiotic sensitive cells in its vicinity under lethal antibiotic concentrations^5,18^.

In bacterial systems, resistance to different antibiotics can exist in separate populations. When these systems are exposed to multiple antibiotics, horizontal transfer of resistance genes may be the only evolutionary pathway for (parts of) the community to survive. As such, our aim was to investigate the transfer of conjugative resistance plasmid to antibiotic susceptible community, if the donor or the recipient, or both were under antibiotic selection. In our experimental setup, the synthetic communities were either sequentially or simultaneously exposed to antibiotics cephalothin and rifampicin (Figure 7A). In practice, *K. pneumoniae, Comamonas* sp. and *Salmonella* Typhimurium strains were co-cultured (N=4) either in 1) cephalothin, 2) cephalothin and rifampicin (simultaneous exposure) or 3) rifampicin. After 30 minutes, *E. coli* donor carrying pEC13 was introduced into the communities, allowing evolutionary rescue to occur. One hour after the introduction of *E. coli* donor, treatments 1 and 3 (i.e. sequential antibiotic exposure treatments), were supplied with rifampicin and cephalothin, respectively. Synthetic communities were cultivated for 3.5 h after which the pEC13 transconjugants were identified via the developed single cell ddPCR method (Figure 7A-B). In treatments 1 and 2, where cephalothin was expected to heavily diminish the initial mock community, the proportion of chimeric 16S rRNA sequences was 15.19% (+/- SEM 0.56%) and 17.27% (+/- SEM 8.05%), respectively. However, in treatment 3, which allowed the initial rifampicin-resistant community to reproduce prior to the second antibiotic treatment, the proportion of chimeras was only marginal (0.42%, +/- SEM 0.40%). These results suggest that high frequency of non-viable cells may have affected the library preparation via the production of chimeric reads, hence, they were excluded from the sequence analysis.

**Figure 7.**
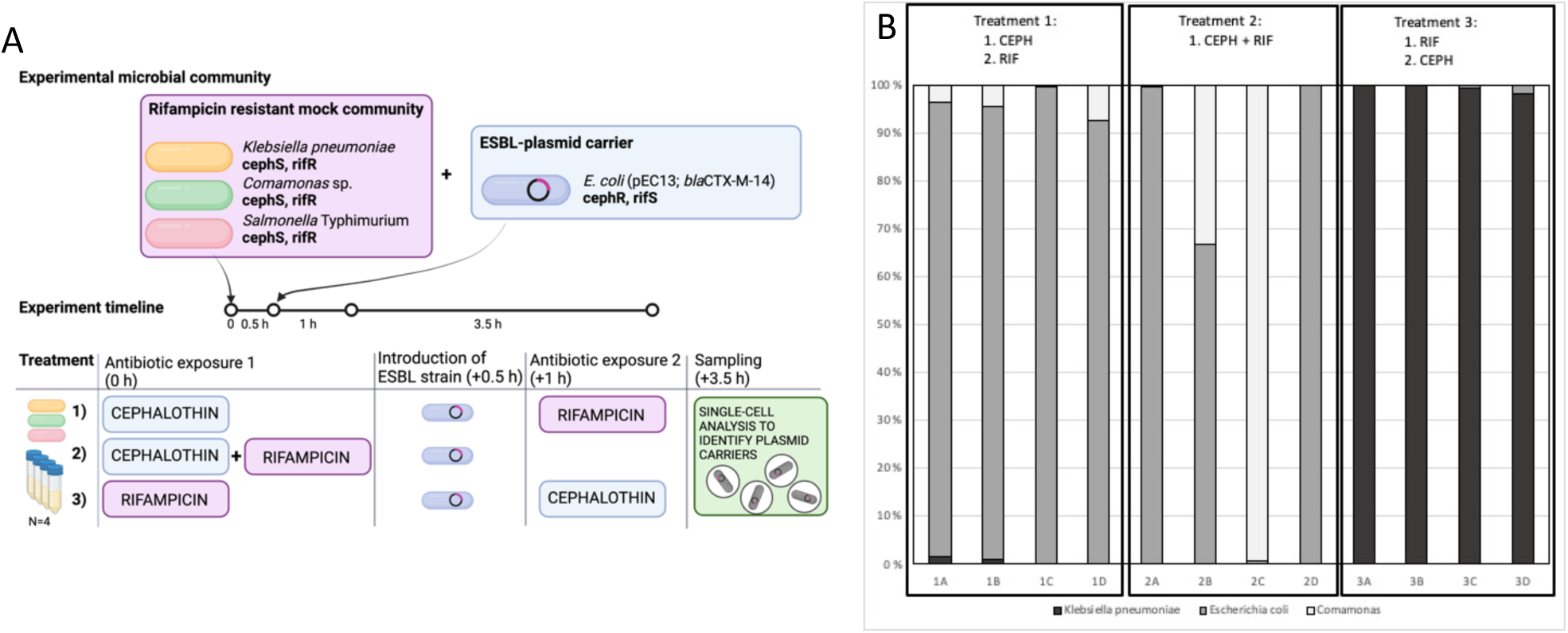
Identifying the ESBL-plasmid carrier population (harboring *bla*CTX-M-14 resistance gene) after sequential or simultaneous antibiotic exposure. A) Rifampicin resistant mock communities consisting of *Klebsiella pneumoniae, Comamonas* sp. and *Salmonella* Typhimurium were exposed to antibiotics in three different treatments (N=4): 1) cephalothin (CEPH), 2) cephalothin and rifampicin (RIF) or 3) rifampicin. After 30 minutes, *Escherichia coli* (pEC13 with *bla*CTX-M-14) was introduced into the communities. The second antibiotic exposure was given after 1 h with 1) rifampicin and 3) cephalothin. Horizontal transfer of pEC13 within the community was determined with single cell ddPCR (*bla*CTX-M-14 and 16S rRNA fusion) after 3.5 h co-culture and is presented as the frequency of plasmid host species **(B)**.

The bacterial community carrying pEC13 was strongly affected by the order in which the antibiotics were introduced into the experimental system (Figure 7B). In treatment 1, in which the community was first exposed to cephalothin, the plasmid carrier population was dominated by the original *E. coli* host. In treatment 3 (first exposed to rifampicin), however, the most abundant plasmid carrier species was *K. pneumoniae*. Simultaneous exposure to both antibiotics led to relatively stochastic behavior of the system, the most abundant plasmid hosts being *E. coli* and *Comamonas* sp. Yet, further pairwise conjugation tests showed that pEC13 conjugation between *E. coli* and *Comamonas* sp. was unlikely to occur (data not shown).

## DISCUSSION

Characterizing microbial communities at the resolution of their true operating units, individual cells, is essential for understanding the functions and biological interactions that take place in complex systems. In this study, we provide a stepwise pipeline that utilizes droplet microfluidics for partitioning bacterial samples into single-cell -containing compartments. The method employs fusion PCR that links a 16S rRNA taxonomic marker with the target functional gene (here an antibiotic resistance gene *bla*CTX-M-14), thus enabling the identification of bacteria carrying the target gene after sequencing the paired amplicons.

Microbial communities, albeit natural or experimental, are supplemented with exDNA that can prevent accurate analysis of genetic material that solely resides within the boundaries of individual viable cells. exDNA does not only originate from lysed cells, but it is also actively released from bacteria because of different functionalities in the microbial system, such as to aid biofilm formation or mediate horizontal gene transfer (see ^19,20^ for review). To exemplify, Carini and colleagues demonstrated that extracellularly located relic DNA may significantly bias the observed microbial diversity in soil samples^21^. This bias, however, was shown to be corrected by treating the soil samples with PMA. PMA is a DNA-intercalating dye that only enters dead cells or cells with compromised membranes and hence it has been used in microbiology to identify the viable bacterial population. The covalent binding of PMA to DNA occurs via photoactivation, preventing the DNA from serving as a template for PCR^22^. While exDNA skews the community analyses when conducted on total microbial DNA, it may also disrupt single-cell methodologies that rely on DNA.

To investigate the prevalence of exDNA in overnight cultures of *E. coli* and *P. fluorescens*, we filtered out the cells and performed 16S rRNA ddPCR for the remaining supernatant. Cell-free supernatant contributed 14-17 % of the total 16S rRNA gene concentration when compared to the corresponding sample where cells were still present. In a preliminary analysis, DNAse treatment was not sufficient in reducing the amplification of exDNA. Hence, the possibility to employ PMA in ddPCR was evaluated. This was done by determining a concentration that did not inhibit the PCR, but effectively blocked the amplification of exDNA in cell-containing and filtrated cell-free samples. The presence of PMA in ddPCR caused a shift in the emitted fluorescence spectrum. However, an accurate categorization of the droplets into positive and negative clusters was still achieved with the 2-dimensional plot of the droplet-emitted fluorescence using two detection channels. As such, potential errors stemming from exDNA can be significantly reduced or eliminated with PMA also in single-cell methods where individual cells and exDNA are trapped within the same compartment.

Production of paired concatemer requires a special primer design that allows amplification and fusion of two physically distant genetic regions. The here-employed protocol includes four primers for amplifying both 16S rRNA and target *bla* regions that are then combined with a fusion primer. Four-primer setup changes the reaction dynamics compared to the three-primer protocol utilized in comparable methods such as epicPCR^12^. In the three-primer protocol, the production of fused concatemer is dependent on the presence of the target gene; in case the trapped bacterial cell does not carry it, neither the 16S rRNA gene nor fusion amplicons are produced. The addition of a fourth primer (in this case, 27F) not only enables the production of 16S rRNA gene fragments of noncarrier cells, but also increases the yield of the fusion product (data not shown). Higher outer primer concentration (compared to inner primers) may enhance the production of fusion concatemer^23^. Thus, we tested several primer ratios and total concentrations in order to achieve the most efficient production of *bla*_16S concatemer. The results showed that the best fusion concatemer production and droplet separation was obtained when outer primer concentrations were several times higher compared to that of inner primers (forward to reverse primer ratios 10:1 for *bla* and 1:5 for 16S rRNA primers, where *bla* forward and 16S rRNA reverse primers acted as outer primers). The application of the here-described system for alternative genetic targets may require optimization of primer concentrations and ratios in order to enhance amplification for better DNA yield and for allowing sufficient separation of negative droplets from positive ones in the subsequent fluorescence analysis.

In this method, the droplets are generated with a microfluidic device that produces equal-sized droplets with a volume of one nanolitre. Bacterial cells are encapsulated into oil-surrounded droplets during droplet generation. The accuracy of ddPCR to quantify DNA concentrations assumes that targets are randomly distributed into droplets of equal size. In the presented approach, individual cells (instead of gene copies) serve as the quantifiable unit. As such, we first assured the applicability of single-cell ddPCR individually for *E. coli, P. fluorescens, K. pneumoniae* and *Salmonella* Typhimurium. To investigate the capability of the method to differentiate *bla*-carriers from non-carriers, *E. coli bla*-carriers were mixed with non-carriers in different ratios and combinations. These mixtures were immediately divided into droplets and the droplets were transferred to a thermal cycler. We employed a prolonged initial denaturation at 95 °C in the ddPCR thermal cycling protocol to ensure cell lysis and accessibility of PCR reagents to the released DNA. As noted above, during PCR, the four primers enable the production of both fusion concatemer from carrier cells and the sole 16S amplicon from non-carriers. As fluorescence is measured after PCR from each droplet one by one, each cell-containing droplet can be detected and hence the ratio of positive to negative droplets can be quantified. The advantage of the QX200 system is that, in addition to its capability to produce uniform droplets, it also inspects the droplet shape and size and excludes abnormal droplets from the analysis. In comparable methods, such as in epicPCR^12^ and OIL-PCR^24^, partitions of various sizes are formed, hence producing compartments with various ratios of PCR reagents to the target DNA. This, in turn, may result in notably differing amounts of amplified DNA that are produced from a single target cell. As such, the here-described method can provide also a more reliable proxy of the number of positive targets in the initial sample, not only their diversity. Further, Poisson distribution is used to select a suitable number of cells to be loaded into a single ddPCR reaction. In other words, a cell concentration that statistically should enclose only a single cell in a droplet while most droplets remain cell-free is used to seed the droplet generation. We confirmed single-cell resolution by parallel single-cell 16S ddPCR to reveal the potential maximum number of positive droplets produced from the sample and proceeded with only those samples that had a target concentration within the confidence interval.

After single-cell ddPCR, the DNA is recovered from all droplets in a single step, thus mixing their contents. At this point, the overall DNA concentration is typically extremely low and needs downstream amplification and barcoding steps to enable amplicon sequencing. The four-primer setup produces single 16S rRNA gene products that can serve as a primer in the subsequent processing and hence may result in false positive host sequences. To prevent cross-binding of fusion amplicons and 16S rRNA products originating from distinct droplets, we used blocking primers in the library preparation steps as described by Spencer and colleagues^12^. An excessive concentration of blocking primers binding to both strands of the 27F region was used to prevent the amplification to start from the 16S rRNA region (*i.e.* in the middle of fusion product, single 16S rRNA gene amplicon or genomic DNA), as in this phase, only fused concatemers are to be enriched. Sequencing of full-length fusion amplicons from mixed cell samples revealed that the vast majority of *bla* host sequences belonged to *E. coli*, which was the only true host carrying the resistance gene. To conclude, our results suggest that the steps for eliminating signals that would potentially cause biased host diversity were sufficient to diminish the false host sequences. However, it should be noted that especially in the case of unknown complex communities, the extremely rare hosts of the target functional gene may be shadowed by the background 16S rRNA gene sequences.

To demonstrate its applicability, we utilized the single-cell ddPCR method for tracing the horizontal transfer of conjugative resistance plasmid pEC13 within a simple synthetic microbial community consisting of cephalothin resistant (rifampicin sensitive) *E. coli* (carrying pEC13 with *bla*CTX-M-14) and rifampicin resistant (cephalothin sensitive) strains of *P. fluorescens, K. pneumoniae* and *Salmonella* Typhimurium. The communities were exposed to three different treatments: 1) cephalothin and after 1 h 30 min to rifampicin, 2) simultaneously to cephalothin and rifampicin or 3) rifampicin and after 1 h 30 min to cephalothin. *E. coli* (pEC13) was introduced 30 min after the first antibiotic exposure. Carriers of *bla*CTX-M-14, and thereby the potential evolutionary rescue via pEC13, were identified with the single-cell ddPCR. When exposed to cephalothin, the susceptible synthetic population was expectedly diminished. Further, after the introduction of *E. coli* with a potential to rescue its accompanying bacteria with the resistance-conferring plasmid, new transconjugant hosts did not seem to appear in the community. In contrast, when rifampicin was administrated as the first antibiotic, the initial resistant population was restored, allowing the spread of rescuing pEC13 plasmid into new hosts before its original host (susceptible to rifampicin) was eliminated from the population. The second antibiotic exposure to cephalothin selects for those bacteria that took in the resistance plasmid and were mainly identified as *K. pneumoniae.* This is in accordance with our previous work where pEC13 was shown to rescue *K. pneumoniae* cells even under lethal cephalothin concentration^18^. The simultaneous exposure to both antibiotics is lethal to all bacterial strains in the community, and therefore assumed not to give enough time for conjugation events. The plasmid host population from this treatment was drastically inconsistent between biological replicates, suggesting that the observed results were likely biased by dead cells dominating the bacterial sample. Moreover, *Comamonas* was not observed to host the pEC13 plasmid, however it was over-represented in some samples, suggesting that certain bacterial behavior, such as secretion of extracellular vesicles, may have significant effects on sequencing results when viable host cells are rare in the community. Nonetheless, these results show that the method can be utilized for identifying the plasmid-carrying sub-community, which itself is shaped by the used antibiotics as well as the resistance phenotypes present in the initial community. Hence, the method has potential for future applications *e.g.,* for studying plasmid-host interactions in more complex microbial communities, such as the gut or soil microbiota – yet it must be acknowledged that different sample types may require further pre-processing steps.

In summary, we have outlined a ddPCR-based protocol that enables both counting the number of bacterial cells as well as providing a phylogenetic assignment of functional genes or extrachromosomal elements (plasmids) to their hosts. Available in many molecular biology laboratories, the ddPCR system offers a valuable droplet microfluidic device for high-throughput genomic research of microbial communities.

## MATERIAL AND METHODS

### Bacterial strains and cultivation

Bacterial strains with and without the target beta-lactam resistance gene were used in the development of the gene tracking method presented in this study. *Escherichia coli* K-12 BL21Gold carrying the conjugative plasmid pEC13 with the target beta-lactamase gene *bla*CTX-M-14^5^ served as the antibiotic resistance gene carrier. *Klebsiella pneumoniae* DSM681, *Salmonella enterica* serovar Typhimurium SL5676, *Pseudomonas fluorescens* (ATCC 13525) and *Comamonas* sp. 4B43 (originating from a human fecal sample) were used as noncarriers. The bacteria were cultivated in Luria Bertani Lennox-broth (LB)^25^, *E. coli, S.* Typhimurium and *K. pneumoniae* at +37°C with 120 or 220 rpm shaking, and *P. fluorescens* at room temperature (RT) with 150 rpm. LB-agar (1%) plates were used for quantifying bacterial densities from liquid cultures. *E. coli, S.* Typhimurium and *K. pneumoniae* plates were incubated overnight at +37°C and *P. fluorescens* 2 days at RT. Media for *E. coli* was supplemented with ampicillin (150 μg/ml) in order to select for pEC13 plasmid.

### ddPCR reactions

The QX200™ Droplet Digital^TM^ PCR system (ddPCR, Bio-Rad Laboratories, Inc.) was used to generate droplets and read the results following the manufacturer’s instructions. The reactions were run in C1000 Touch™ Thermal Cycler (Bio-Rad Laboratories, Inc) with 96 Deep-Well Reaction Module. The ddPCR reactions were kept overnight at 12 °C before reading the droplets. The results were obtained from channels 1 (FAM) and 2 (HEX/VIC) and analysed with QuantaSoft^TM^ Analysis Pro 1.0 (Bio-Rad Laboratories, Inc). All ddPCR reactions were prepared with QX200™ ddPCR™ EvaGreen® Supermix (2x).

### 16S rRNA gene amplification

The amplification of the 16S rRNA gene was done using universal primers 27F and 338R (the primers used in this study are listed in Table 1). After initial optimization of the 16S rRNA primers for ddPCR (see Supplementary Figure S2 for more information), the reactions were prepared using 1x ddPCR supermix and 100 nM of each primer. The amplification conditions consisted of initial incubation at 95 °C for 10 minutes, followed by 40 cycles of denaturation at 95 °C for 60 seconds, annealing at 55°C for 60 seconds and elongation at 72 °C for 2 minutes, and a final step for fluorescence stabilization with incubation at 4 °C for 5 minutes followed by incubation at 90 °C for 5 minutes. The ramp rate between steps was 2 °C/sec until the last elongation incubation, and 1 °C /sec for the fluorescence stabilization. Additionally to the fluorescence analysis by the BioRad droplet reader, a subset of droplets was also imaged under a fluoresence microscope (Zeiss Axio Vert.A1, with Leica MC170HD camera and Zeiss Filter set 09 including Excitation Bandpass Filter 450-490 nm, Emission Longpass Filter 515 nm and Beamsplitter FT 510) to manually assess the difference in fluorescence (Supplementary Figure S4C-D). The produced droplets and the delivery of viable bacterial cells into droplets were inspected under a bright field microscope (Zeiss Axio Vert.A1, with Leica MC170HD camera) (Supplementary Figure S4A-B).

**Table 1.**
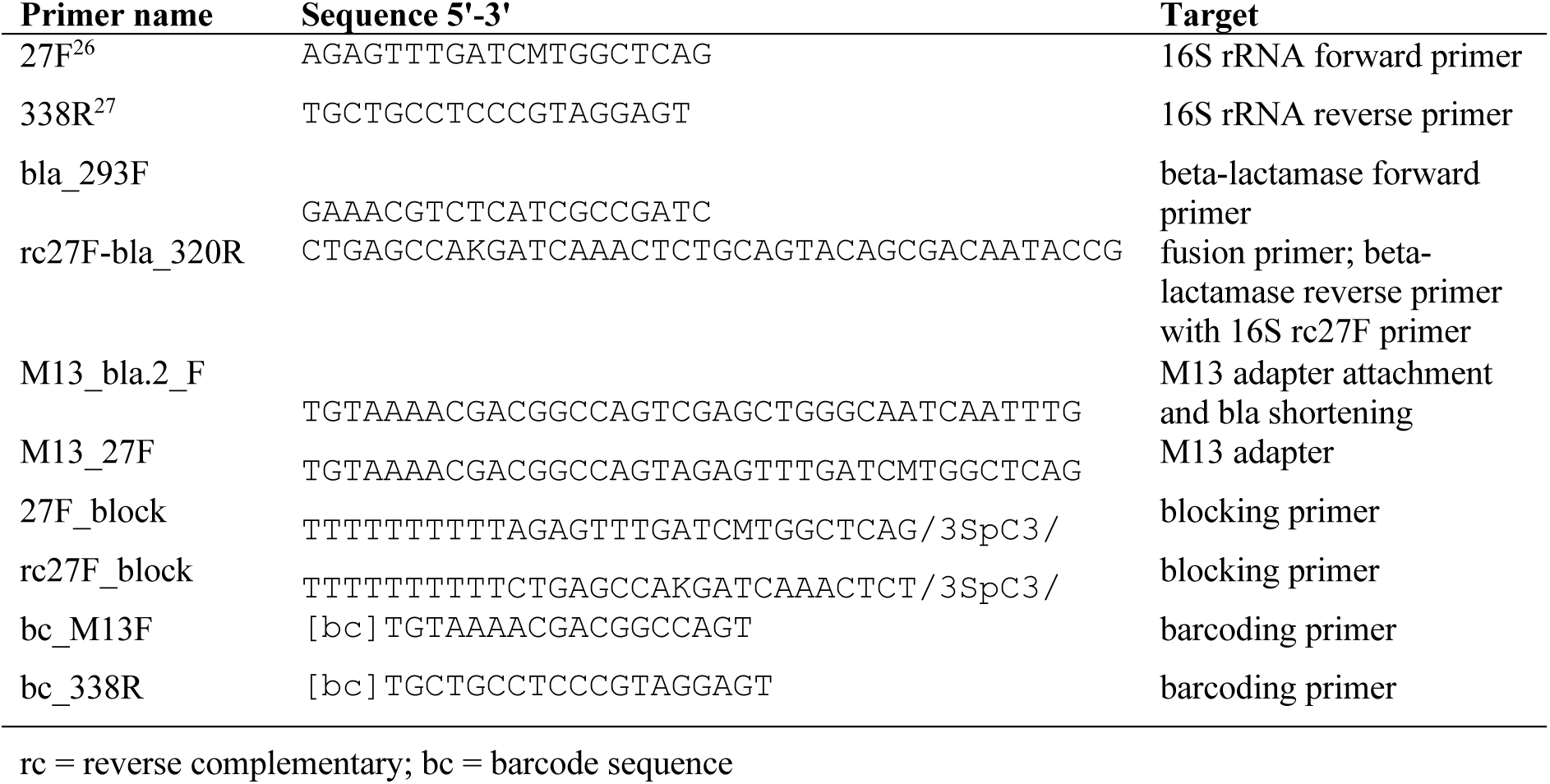
Primers used in this study.

### Propidium monoazide (PMA) for sample treatment

Initially, the maximum non-inhibitory concentration (MNIC) of propidium monoazide (PMA) in ddPCR reaction was tested. PMA (PMAxx^TM^, Biotium) was diluted in PCR grade water to 10 µM of final concentration and photoactivated. The treatment consisted of initial incubation in the dark for 10 minutes, followed by incubation for 10 minutes under blue LED light (460 nm). The solution was prepared in 500 µl thin-walled polypropylene Qubit^TM^ assay tubes (Invitrogen^TM^) with final volume of 200 µl. Samples were kept on ice layer and mixed occasionally during the treatment by inverting the tube. Lower concentrations were prepared from the treated PMA solution and used in ddPCR, with final concentration ranging from 0.125 to 2.5 µM. *E. coli* DNA digested with FastDigest HindIII and NotI (Thermo Fisher Scientific, MA, USA) was used as a template in ddPCR reactions optimized for 16S rRNA. The ddPCR was prepared using 100 nM of each universal primer (27F and 338R) for 16S rRNA amplification. The amplification program was carried out using 5 minutes of initial incubation at 95 °C, and 40 cycles of denaturation at 95 °C for 30 seconds, annealing at 55 °C for 60 seconds and elongation at 72 °C for 2 minutes. The final fluorescence stabilization and ramp rate were the same as mentioned above.

Next, cell samples prepared from *E. coli* and *P. fluorescens* cultures were treated with PMA and used as template in ddPCR. Bacterial suspensions were freshly prepared for every new assay by diluting an overnight culture up to 1000-fold in 1x phosphate buffer solution (PBS). The putative inhibitory effect of PBS (from concentration ranging from 0.015625x to 0.25x of PBS) on ddPCR efficiency was determined during the initial optimization of reaction conditions for 16S rRNA primers (Supplementary Figure S1). After determining the maximum non-inhibitory PBS concentration, it was always kept under 0.0625x in the ddPCR reactions. The filtrate was obtained by individually centrifuging 250 µl of each dilution at 5000 × *g* for 3 minutes through a 0.2 µm spin filter (Centrifugal Filter, Modified Nylon 0.2 μm, VWR). Both sample types, cell suspension and filtrate, were treated with 10 µM of PMA as described above. After PMA treatment, the cells were diluted in PCR grade water and used as template in ddPCR reaction using 16S rRNA primers. The same reaction conditions used for the MNIC assay were used here. The cell density of the samples was determined by serial dilution and plate cultivation of cell suspensions, before and/or after PMA treatment.

### One-step fusion PCR for concatemer production

Specific primers were designed to detect the pEC13 plasmid and prompt the bla_16S concatemer formation. The plasmid sequence was retrieved from the GeneBank database under accession number KU932024.1. The class A beta-lactamase gene (*bla*CTX-M-14) present in the plasmid was the target sequence chosen and here referred to as *bla*. Primers were designed using the online Primer Designer™ Tool (Thermo Fischer Scientific). To amplify a 101-bp *bla* fragment, 293F was used as a forward primer together with a modified reverse primer rc27F-bla_320R (Table 1). It contains *bla* reverse primer part (320R) and 5’-overhang encoding the reverse complement sequence of 16S rRNA forward primer 27F. A fusion primer enables the production of the 450-bp concatemer via the attachment of the amplified *bla* fragment on its 27F-overhang to 16S rRNA fragment (350 bp PCR product amplified by primers 27F and 338R) in the same reaction (Figure 2). Initially, both primer pairs (for amplifying *bla* and 16S fragments) were screened separately for forward to reverse ratio and primer pair concentration in ddPCR (Supplementary Figure S3). Reactions were prepared with up to a 100-fold difference between forward and reverse primer molarity, and total concentration ranging from 100 to 400 nM. After the initial screening, both primer pairs were used in the same reaction to produce the concatemer. The total concentration of 16S rRNA primers was fixed to 225 nM and the ratio of forward to reverse to 1:5. *bla* primers were used in concentrations ranging from 100 to 300 nM and ratios from 10:1 to 30:1 (Supplementary Figure S3C-E). Finally, the primer ratios and concentrations were defined for single-cell experiments. The further reactions for concatemer production were prepared with 225 nM and a ratio of 1:5 (forward to reverse) of 16S rRNA primers, and 200 nM and 10:1 (forward:reverse) of *bla* primers.

### Single bacterial cells in droplets

To test whether single bacterial cells are suitable to be used as templates in ddPCR, overnight cultures of *E. coli, P. fluorescens, K. pneumoniae,* and *S.* Typhimurium were diluted to reach a cell density of approximately 1×10^7^ cfu/ml (cfu; colony forming units). Bacterial dilutions were PMA-treated as described above with the final PMA concentration of 10 µM. A dilution series of 1:2 was prepared (in PCR grade water) from the PMA-treated cell samples and used as a template in ddPCR. The bacterial cell suspension always constituted ¼ of the total ddPCR reaction volume. The maximum detectable cell number was determined by using 16S rRNA primers 27F and 338R. The corresponding bacterial cell density of the samples was also determined by dilution series and plating on LB plates. The protocol optimized for the amplification of the 16S rRNA gene in ddPCR and specified in the section above was used.

### Detection of beta-lactamase host bacteria from mixed samples with single-cell ddPCR

Overnight cultures of *E. coli, K. pneumoniae,* and *S.* Typhimurium were diluted for PMA-treatment and a 1:2 dilution series was prepared as described above. Diluted bacterial samples containing either one bacterial strain or a mix of two strains: beta-lactamase -carrying *E. coli* and a non-carrier strain *K. pneumoniae* or *S.* Typhimurium was used as template in ddPCR. Both 16S rRNA (amplified with 27F and 338R) and concatemer bla_16S (amplified with four primers; 293F, rc27F-320R, 27F and 338R) were amplified from the same samples in separate reactions. Two replicates were prepared for reading the droplets from the 16S rRNA reaction, and three replicates were prepared for the one-step fusion reaction to be saved and used in the following procedures. The ratio of positive:negative droplets was determined with the Droplet Reader and the samples containing a sufficient number of positive droplets according to 16S rRNA amplification (and therefore verified to contain the cell density required for single cell analysis) were proceeded to DNA recovery. The amplification conditions were the same as in the protocol optimized for 16S rRNA gene amplification in ddPCR (described in the specific section above).

### DNA recovery from droplets and library preparation for sequencing

The amplified *bla*_16S concatemers were recovered from droplets and enriched before preparing the sequencing libraries. The recovery was done by freeze-thawing the ddPCR emulsion using liquid nitrogen (LN2), a method we described recently^17^. After freeze-thawing, phase separation was achieved, and the amplicon-containing phase was collected for further processing. The recovered concatemers were purified with the Agencourt AMPure XP purification system (Beckman Coulter, Brea, CA) using a 0.9x ratio.

Sample enrichment was performed before library preparation by reamplifying the recovered *bla*_16S amplicons with primers M13_bla.2_F and 338R (Table 1). The samples were then barcoded with bc_M13 and bc_338R primers, both extended with unique 5’-barcode sequences (bc). Both enrichment and barcoding qPCR reactions were prepared with 1x Maxima SYBR Green/Fluorescein qPCR Master Mix (ThermoScientific^TM^) in a final volume of 25 µl. To prevent the undesired pairing of bla-27F overhang with other 16S rRNA fragments that were present in the sample, the reactions included blocking primers block_27F and block_27R, both in a final concentration of 1 µM. The product of barcoding qPCR was cleaned by gel extraction using NucleoSpin Gel and PCR clean-up kit (MACHEREY-NAGEL, GmbH & Co, KG, Germany), after 1% AGE (agarose gel electrophoresis) run for 1 hour at 100 V. The fragment was further analysed with High Sensitivity D1000 ScreenTape assay in 2200 TapeStation System (Agilent Technologies, Inc, CA, USA). PCR conditions for enrichment and barcoding reactions were as follows: the amplification run started with pre-incubation at 95 °C for 5 minutes, followed by 15 cycles of denaturation at 95 °C for 30 seconds, annealing at 53 °C for 30 seconds, and elongation at 72 °C for 30 seconds (with plate read after each cycle). After the run, the product was analysed in a 1.5 % AGE run for 1 hour at 110V. The barcoded and purified *bla*_16S amplicons were proceeded to library preparation with NEBNext® Ultra™ II DNA Library Prep Kit and sequenced with Illumina NovaSeq 6000 platform using SP flowcell (PE250). Further sequence analysis of 250-bp paired-end read data was run with CLC Genomics Workbench 12 (Qiagen Bioinformatics). The obtained reads were trimmed with *bla* and 16S rRNA primer sequences bla_293F, bla_320R, 27F and 338R to confirm that they contain both parts of the concatemer. The genetic region between 27F and 338R was extracted and used for *de novo* OTU clustering with a 97 % similarity threshold. Rare OTUs (abundance <100 reads) were excluded from the analysis. The reference sequence of each OTU was exposed to BLAST search, using the 16S Ribosomal RNA sequences database. Each OTU was identified according to the BLAST results. Finally, the abundance of identified bacterial hosts carrying *bla*CTX-M-14 present in each sample was determined.

### Horizontal transfer of beta-lactamase under lethal antibiotic exposure

Rifampicin-resistant strains of *K. pneumoniae, S.* Typhimurium and *Comamonas* sp. (bacterial isolate originating from the human fecal sample as part of gut bacterial strain library collected under research permit T300/2020 for Hospital District of Southwest Finland) were used to construct a synthetic bacterial community. 5 µl of each bacterial culture, cultivated overnight in LB medium without antibiotics, were added to 5 ml of fresh LB medium containing 1) cephalothin 2) rifampicin and cephalothin or 3) rifampicin, with four replicates per treatment. The bacterial communities were incubated at +37 °C for 30 min with 120 rpm agitation, after which *E. coli* harboring conjugative plasmid pEC13 with *bla*CTX-M-14 (conferring resistance to cephalothin) was introduced by adding 5 µl of overnight grown culture into each bacterial community. After 1 h incubation, a second antibiotic pulse was given to treatment 1 with rifampicin and treatment 3 with cephalothin, resulting in the same final antibiotic concentration in all the treatments (50 μg/ml of both cephalothin and rifampicin). The bacterial communities were incubated for 3.5 h after which one cell sample from each community was taken for single cell analysis for tracing the *bla*CTX-M-14 carrier cells.

Cell samples from each community were PMA-treated and diluted as described earlier. *bla*_16S concatemers were produced by single-cell ddPCR. Each reaction was run in two replicates. One reaction was read with Droplet Reader to ensure cell density and single-cell resolution. The other replicate was used for DNA recovery with LN2. Recovered fusion amplicons were first purified with QIAquick PCR purification kit (Qiagen) and proceeded to library preparation first with re-amplification PCR (PCR conditions: 95 °C 5 min, followed by 30 cycles of denaturation at 95 °C for 30 seconds, annealing at 53 °C for 30 seconds, and elongation at 72 °C for 30 seconds, with plate read after each cycle) followed by PCR purification (sparQ PureMag Beads, Quantabio) and inspecting the correct amplicon size with AGE (1.5%, 30 min, 120 V). Purified fusion amplicons were barcoded with both 5’ and 3’ extensions in barcoding PCR (PCR conditions as described above but with 9 cycles). Barcoded amplicons were extracted from agarose gel (1.5%, 1 h, 100 V) by using NucleoSpin® Gel and PCR Cleanup XS kit (Macherey-Nagel) and the amplicon size and DNA concentration were checked with High Sensitivity D1000 ScreenTape assay in 2200 TapeStation System (Agilent Technologies, Inc, CA, USA). Sequencing libraries were prepared with NEBNext® Ultra™ II DNA Library Prep Kit and the samples were sequenced with Illumina NovaSeq 6000 platform using SP flowcell (PE250).

Further sequence analysis of 250-bp paired-end read data was run with CLC Genomics Workbench 21 (Qiagen Bioinformatics) and Microbial Genomics module. Paired reads were first merged and trimmed by quality (0.05 as threshold). Reads were then trimmed with forward and reverse primer sequences of *bla*CTX-M-14 to ensure they contain the complete sequence of the target gene, and then with 16S forward primer. Single reads and merged read sequences not covering the complete fusion amplicon length were discarded from the analysis. Finally, the remaining sequence between 16S rRNA forward (27F) and reverse (338R) primer binding sites was used to determine the carriers of the beta-lactamase gene. Only the reads that contained all the primer sequences and that had a length of 300-315 bp (after trimming) were taken to the final analysis. A local 16S database was constructed to represent the synthetic community (consisting of *K. pneumoniae, S.* Typhimurium, *Comamonas* sp. and *E. coli* 16S rRNA gene sequences) and used for reference-based OTU clustering with 99% similarity, allowing the creation of new OTUs (99% taxonomy similarity). OTUs with a total abundance lower than 4000 or recognized as chimeras were left out from the final analysis. The subpopulation carrying *bla*CTX-M-14 was assessed as OTU frequency (OTU abundance/total OTU abundance).

## Acknowledgements

This study was funded by the Academy of Finland grants 347531 to M.J., 323063 to M.T. and 322204 to R.P., ERC Consolidator Grant (615146) to M.T., Finnish Cultural Foundation grant (00210277) to L.A.L.D. and Jane and Aatos Erkko Foundation to M.J.

## Supplementary Data

**Supplementary Figure S1.**
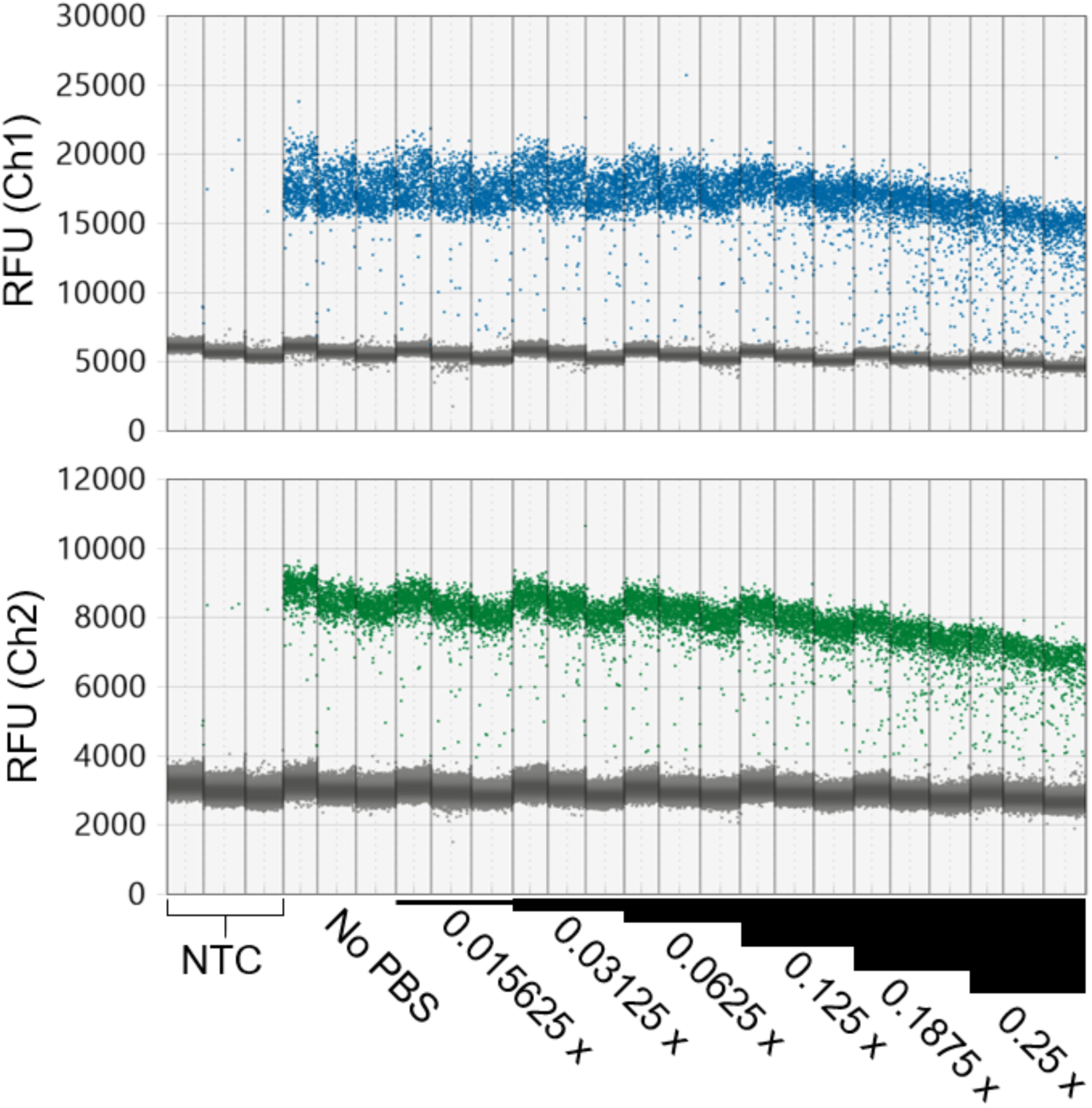
Inhibitory effect of PBS in ddPCR reaction. PBS dilutions were added to ddPCR reactions to detect inhibition of amplification. The amplification of 16S rRNA using the ddPCR reaction optimized in this study was used to demonstrate the effect of PBS in ddPCR. *E. coli* DNA was used as a template in the reactions and PBS concentrations ranging from 0.0156x to 0.25x were tested. After the amplification, the RFU (relative fluorescence units) of each droplet was analysed from channels 1 (Ch1; FAM) and 2 (Ch2; HEX/VIC). The amplitude of positive clusters reduced with increasing concentrations of PBS, and more droplets were observed to form the rain effect in reactions with 0.125-0.25x of PBS. These effects could be seen from both channels. PBS concentration of 0.0625x was set as the maximum non-inhibitory concentration for ddPCR reactions.

**Supplementary Figure S2.**
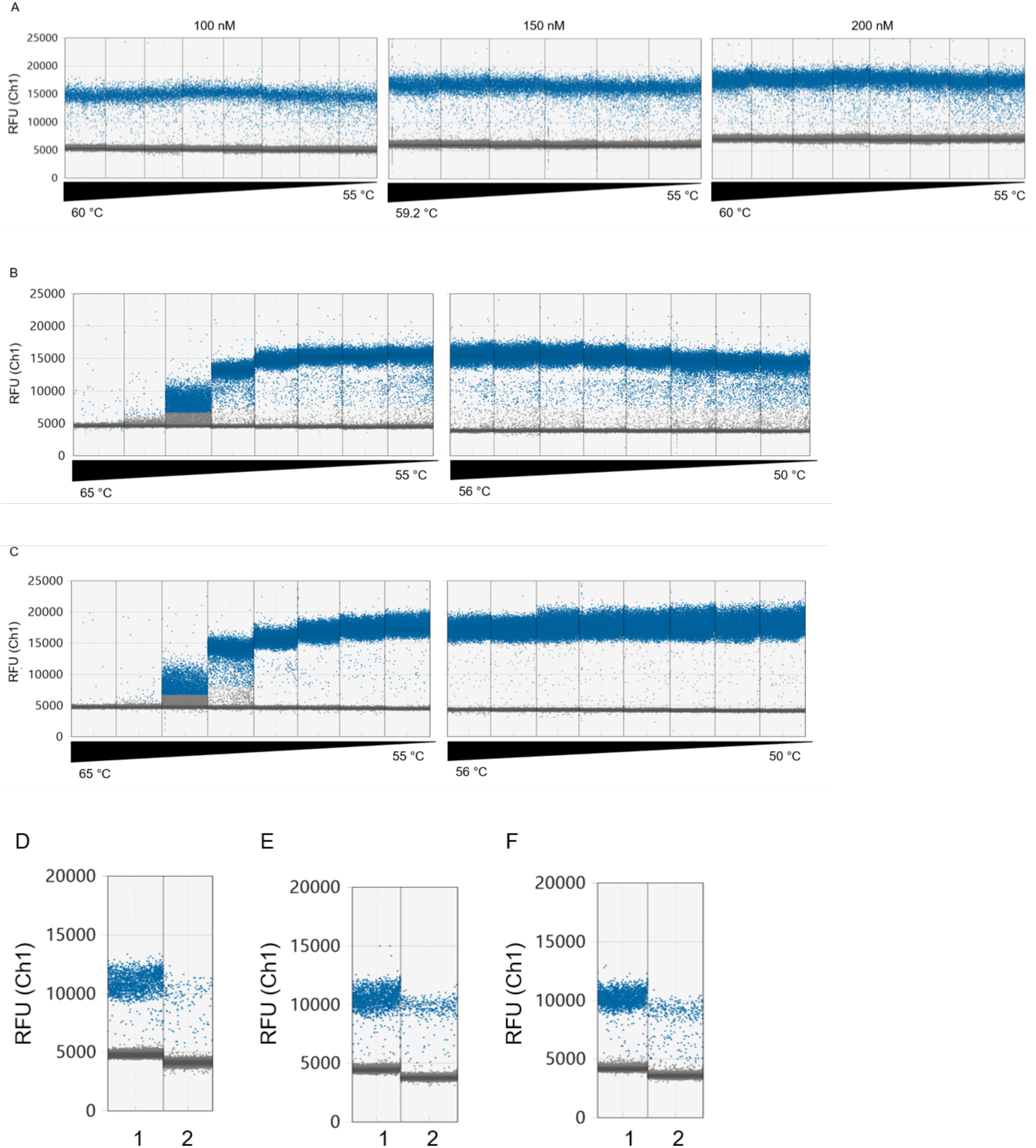
Optimization reactions for 16S rRNA amplification in ddPCR using universal primers. The amplification of 16S rRNA using universal primers 27F and 338R was optimized for ddPCR by testing primer concentration, annealing temperature, duration of initial incubation and the cycling steps. Initially, three primer concentrations (100 nM, 150 nM, 200 nM) were tested in a cycling protocol using an annealing temperature gradient between 60 and 55 °C (A). Heavy “rain effect”, *i.e.* droplets’ fluorescence not clustering with positive nor negative droplets, was observed in higher primer concentrations and lower annealing temperature. Cycling protocol with two (denaturation and annealing) (B) or three steps (denaturation, annealing, and elongation) (C) were tested *versus* annealing temperature gradients from 65 to 55 °C and 56 to 50 °C. The rain effect was more pronounced in two-step cycles and lower temperatures. Duration of initial denaturation and cyclic denaturation steps were tested using *P. fluorescens* (1) and *E. coli* (2) cell suspensions to optimize the detection of single cells. Initial denaturation at 95 °C for 5 minutes (D), 10 minutes (E) and 95 °C for 5 minutes with a longer cyclic denaturation (1 minute at 95 ° C) (F). The positive droplets formed a narrower cluster in reactions using either longer initial (E) or longer cyclic denaturation (F), when compared with the standard protocol (D). The rain effect decreased in longer initial denaturation. In combination with these results, the following settings were adopted for further assays: long initial incubation (10 minutes at 95 °C); three-step cycling assay with long denaturation (1 minute at 95 °C), annealing at 55 °C (1 minute), and elongation at 72 °C for 2 minutes.

**Supplementary Figure S3.**
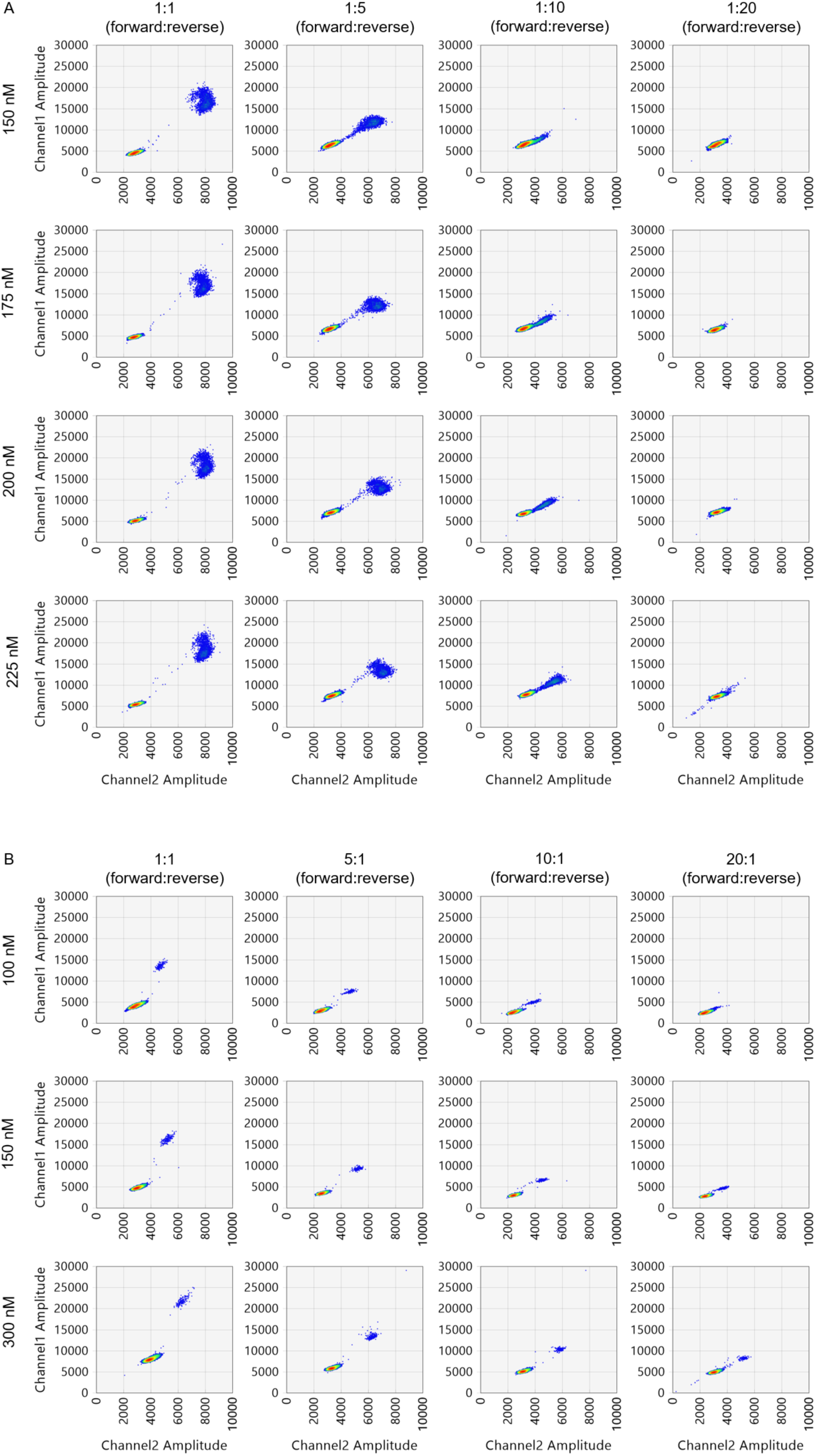
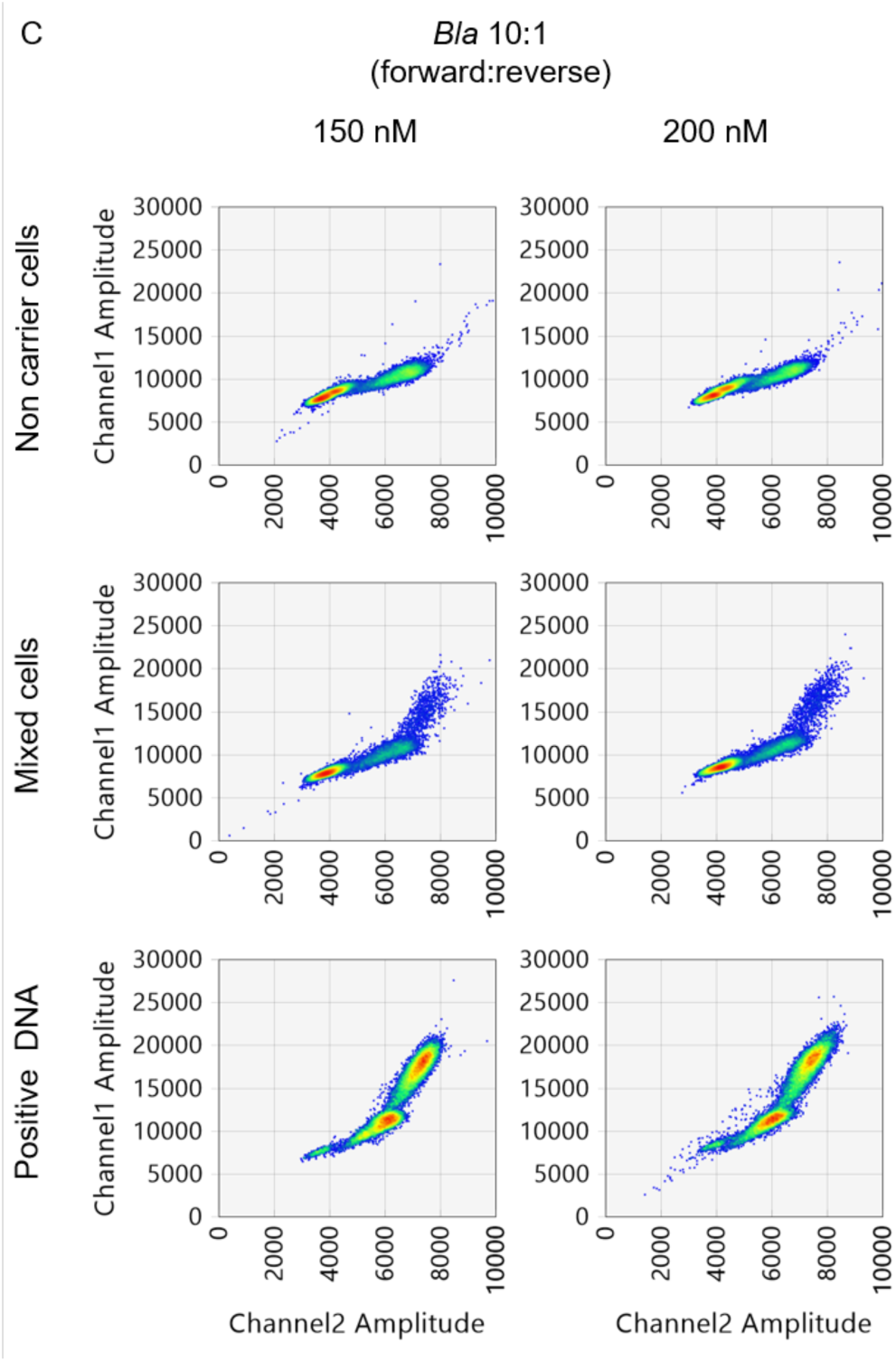
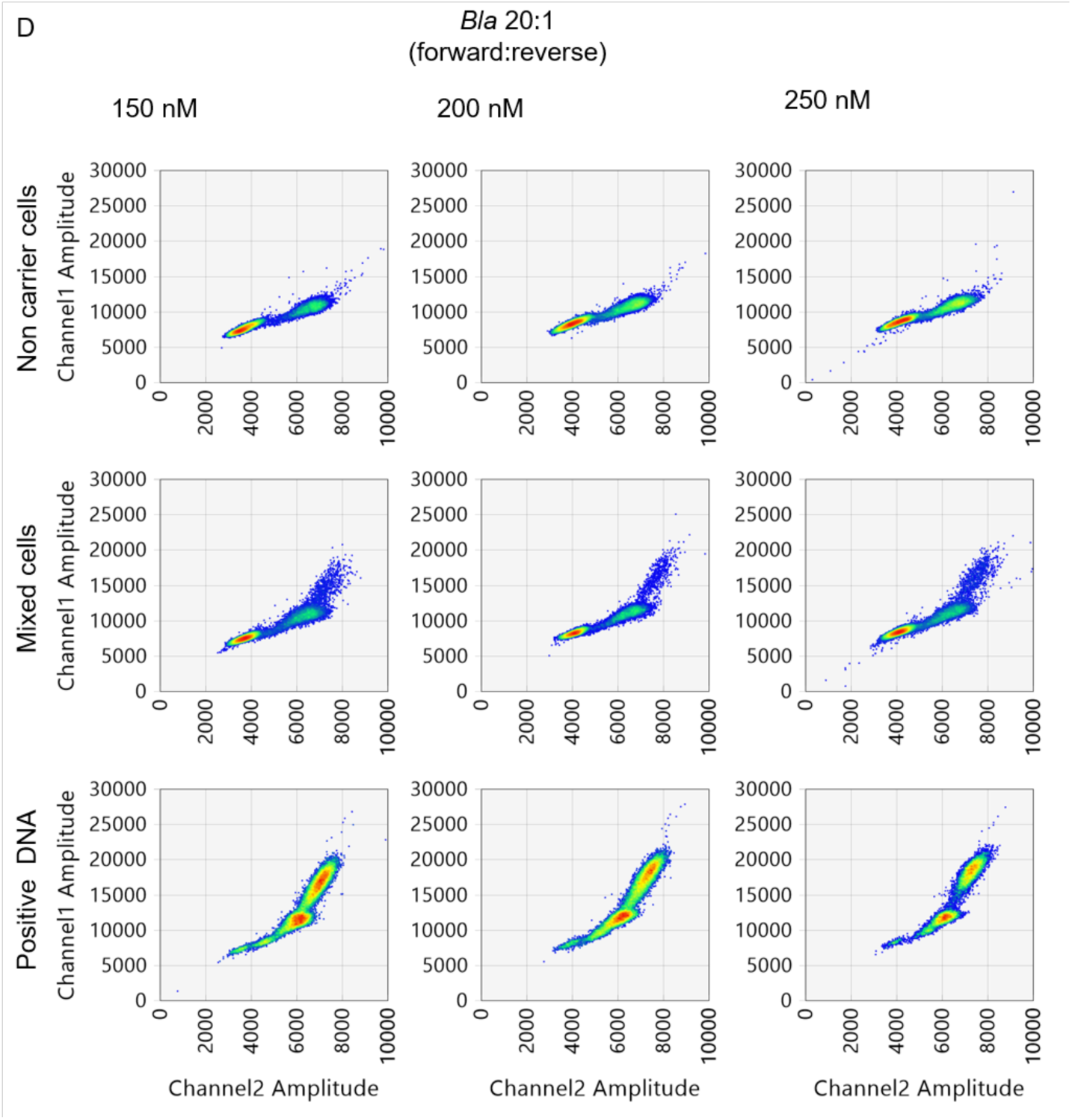
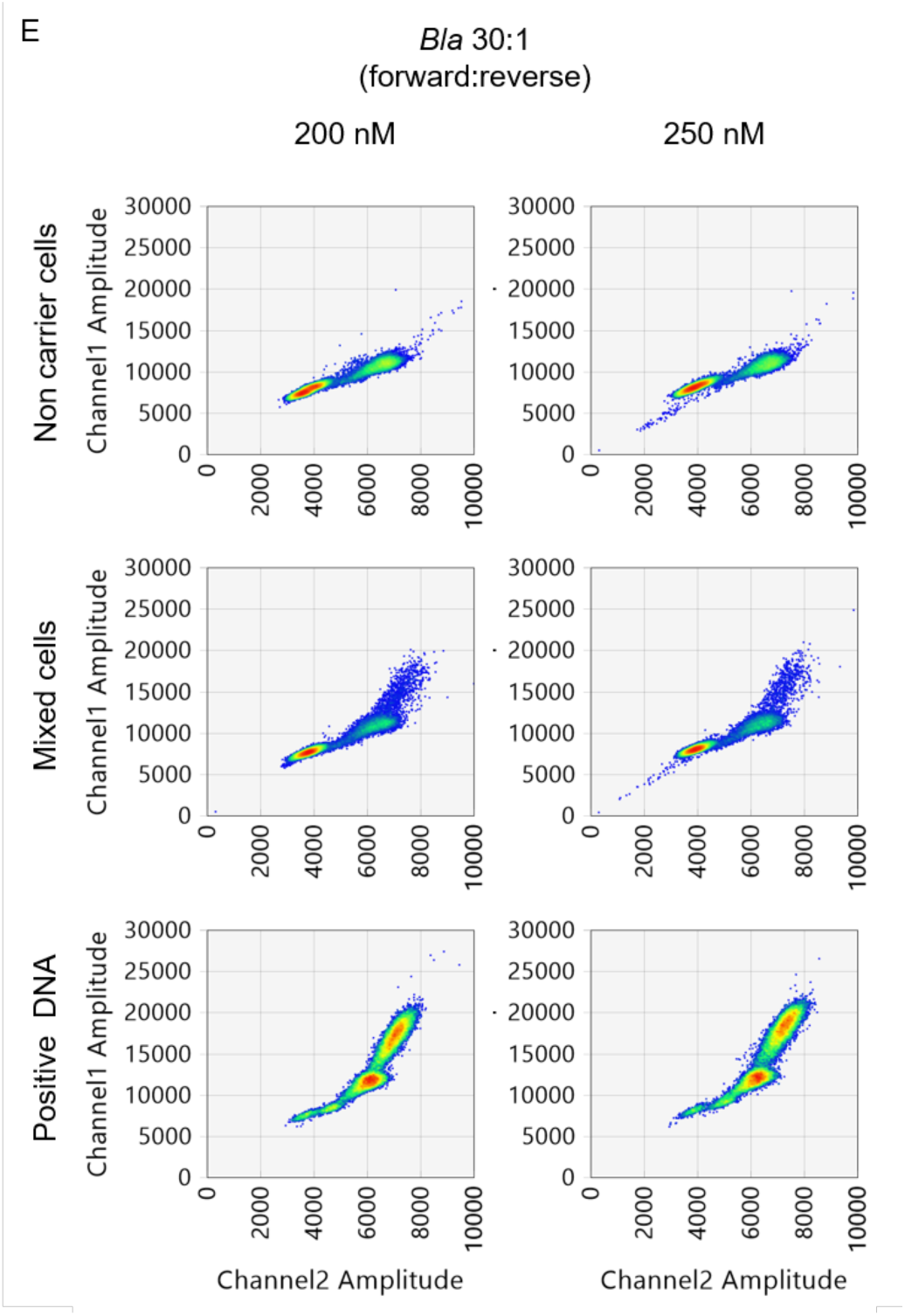
One-step fusion ddPCR optimization. The fusion reaction to generate the concatemer of 16S rRNA and *bla* was optimized in ddPCR by testing asymmetric molarity of primers in pairs, and varying concentration of each pair. The ratio and concentration were first screened separately for each pair, using *E. coli* DNA as a template. The fluorescence of each droplet was analysed, and the formation of distinct clusters for positive and negative droplets was checked for each sample. 16S rRNA primers were tested with concentrations varying from 150 to 225 nM and ratios 1:1 to 1:20 (forward:reverse) (A). Primers for *bla* were tested with concentrations of 100, 150 and 300 nM and ratio up to 20:1 (forward:reverse) (B). After being tested in separate reactions, both primer pairs were used in the same ddPCR to produce bla_16S concatemer. Bacterial cell suspension of *E. coli* carrying *bla, Pseudomonas fluorescens* (non-carrier strain), and DNA was used as template in these reactions. The reactions were prepared with a fixed (total) concentration (225 nM) and ratio of 1:5 (forward:reverse) of 16S rRNA primers. Selected concentrations of 150, 200 and 250 nM of *bla* primers were used in one-step fusion ddPCR assay with forward:reverse ratios of 10:1 (C), 20:1 (D) and 30:1 (E).

**Supplementary Figure S4.**
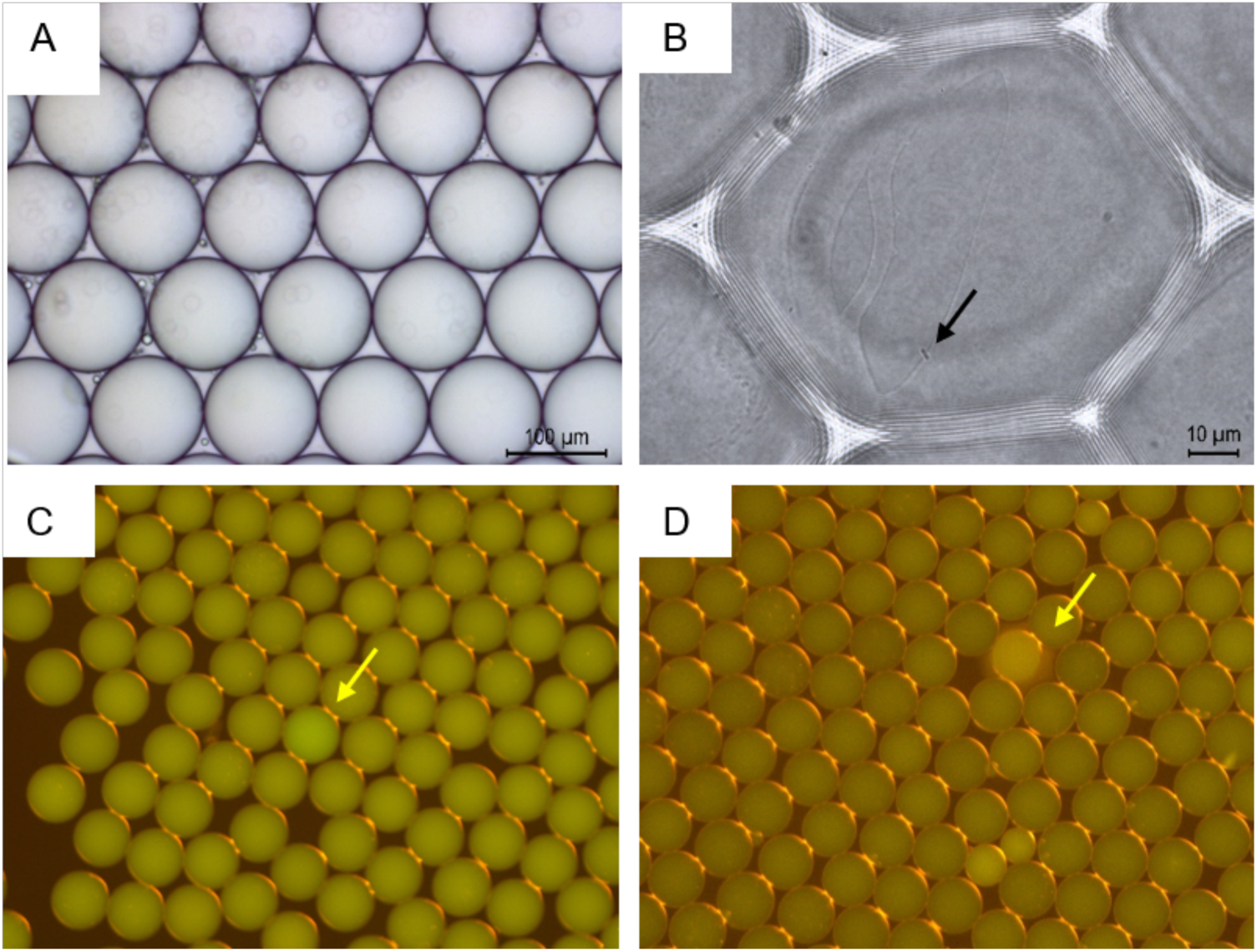
Droplet partitions generated for single-cell ddPCR. Droplets were generated with ddPCR system, single *E. coli* cells were delivered into droplets and imaged using bright field microscopy (Zeiss Axio Vert.A1, with Leica MC170HD camera) (A-B). The black arrow in panel B points to a bacterial cell within a ddPCR partition. Droplets prepared using *E. coli* DNA as a template were subjected to single-cell 16S rRNA ddPCR and imaged using fluorescence microscope (Zeiss Axio Vert.A1, with Leica MC170HD camera and Zeiss Filter set 09 including Excitation Bandpass Filter 450-490 nm, Emission Longpass Filter 515 nm and Beamsplitter FT 510) (C-D). The yellow arrows point to droplets with the higher fluorescence emission.

